# Patterns of genetic variation in a prairie wildflower, *Silphium integrifolium*, suggest a non-prairie origin and locally adaptive variation

**DOI:** 10.1101/2020.06.25.171272

**Authors:** Andrew R. Raduski, Adam Herman, Cloe Pogoda, Kevin M. Dorn, David L. Van Tassel, Nolan Kane, Yaniv Brandvain

## Abstract

**Premise:** Understanding the relationship between genetic structure and geography provides information about a species’ evolutionary history and can be useful to breeders interested in de novo domestication. The North American prairie is especially interesting because of its relatively recent origin and subsequent dramatic fragmentation and degradation. *Silphium integrifolium* is an iconic perennial American prairie wildflower targeted for domestication as an oilseed crop. Germplasm in the existing breeding program is derived from accessions collected in restricted geographic regions. We present the first application of population genetic data in this species to address the following goals (1) improve breeding programs by characterizing genetic structure and (2) identify the species geographic origin and potential targets and drivers of selection during range expansion.

**Methods:** We developed a reference transcriptome as a genotyping reference for samples from throughout the species range. Population genetic analyses were used to describe the distribution of genetic variation and demographic modeling was used to characterize potential processes that shaped variation. Outlier scans for selection and associations with environmental variables were used to identify loci linked to putative targets and drivers of selection.

**Key results:** Genetic variation partitions samples into three geographic clusters. Patterns of variation and demographic modeling suggest that the species origin is in the American southeast. Breeding program accessions are from the region with lowest observed genetic variation.

**Conclusions:** This iconic prairie species did not originate within the modern prairie. Breeding programs can be improved by including accessions from outside of the germplasm founding region, which has relatively little variation. The geographic structuring coupled with the identified targets and drivers of adaptation can guide collecting efforts towards populations with beneficial agronomic traits.

## Introduction

The North American prairie was once the largest vegetative region on the continent. It has served as a cultural identity reference point as well as the subject of many grassland ecological studies (Weaver 1954; Lauenroth 1979; Matthews 1988; Wilson and Hartnett 1998; Samson et al. 2004; Knapp et al. 2008). The overwhelming majority of the prairie has since been altered by human disturbance with some estimates of range decline since European settlement as high as 99.9% (Samson and Knopf 1994). Conservation efforts of this endangered habitat have focused on protection of virgin prairie fragments and restoration of altered landscapes by human directed reseeding of prairie plants. The latter approach is far more common, in part, because of the scarcity of available virgin prairie. Understanding the forces that influence genetic structuring of species is a fundamental goal of population genetics and it is not yet well understood how the very recent prairie decline affects patterns of natural variation established prior to the onset of decline, especially in long-lived perennial plants. Having knowledge about how genetic variation is distributed across natural landscapes can be helpful towards deciding which virgin prairies to prioritize or which geographic ecotypes of plants species to plant in prairie restorations.

Surprisingly little is known about the evolutionary history of common North American prairie forbs from population genetic data, and it is not currently known if there are general patterns across species of directional range expansion following glaciation or locations of geographic sources of ancestral variation. Changes in species composition have drastically altered the biome as C4 grasses became dominant on the landscape over the past few million years (Fox and Koch 2003, 2004). Data from fossil pollen deposits have suggested that now common prairie wildflower families, such as the Asteraceae, were likely not plentiful on the prairie landscape at the time that C4 grasses became dominant (Williams et al. 2004). Understanding the evolutionary and demographic history of how a species has come to inhabit its range can allow us to better understand the present relationship between a species’s genetic variation and geography. Currently, the extent to which recent habitat fragmentation affects genetic signatures of past range expansion and how a species life history affects the pattern is not well understood. Perennial plants may be buffered more to a degree than annuals from genetic diversity losses brought on by habitat fragmentation.

Prairie perennial plants have become the inspirations and models for neo-domestication efforts to develop sustainable agriculture. Roots of perennial plants in the soil protect against soil erosion, promote a diverse soil microbiome, and act as a carbon sequestration resource. Species of the genus *Silphium* (Asteraceae) are deep-rooted, long-lived, perennial wildflowers with flowers pollinated by bees and seeds dispersed by wind that are found throughout the United States, east of the Rocky Mountains. Common *Silphium* species are best known for being charismatic flora of the North American prairie and their ability to tolerate extreme abiotic stresses (Weaver et al. 1935; Leopold 1968). Recently there have been efforts to better incorporate the ecological benefits of *Silphium* species into improving habitat restoration and agricultural practices (Van Tassel et al. 2017). Because of these benefits, combined with high biomass accumulation requiring relatively few inputs, species of *Silphium* show promise for forage and biogas production (Ustak and Munoz 2018; Wever et al. 2019) or as a perennial oil seed crop (Van Tassel et al. 2017). Additionally, the incorporation of native wildflowers in prairie restoration projects or as buffer strips among agriculture fields are thought to improve pollinator abundance (Isaacs et al. 2009). *Silphium* species are known to attract beneficial insects (Fiedler and Landis 2007) and are currently being evaluated for such conservation practices because they are large plants, competitive with weeds after established, have potential for economic return (seed or bioenergy) and breeding programs may improve seedling vigor compared to other wildflowers (Butters et al. 2019). Despite increased interests across applied settings, relatively little is known about patterns in the distribution of genetic variation in long-lived perennial prairie genera such as *Silphium*, or how recent habitat fragmentation and degradation have affected levels of variation in natural populations.

As there has been a recent movement to domesticate *S. integrifolium* and *S. perfoliatum*, understanding the underlying extent and distribution of genomic variation within and among populations will help us better understand how recent habitat fragmentation has affected the genetic signature of range expansion and also support ongoing applied *Silphium* research. Breeding trials to develop *S. perfoliatum* as a potential cover crop began in the 1950’s and continue today with additional effort to develop it as a biogas resource (Stanford 1990; Wever et al. 2019). Most recently, *S. integrifolium* has been targeted for development as an oilseed crop, in part, because of the similar fatty acid composition profile in its seeds to sunflower seeds (Reinert et al. 2019). Accessions used in *S. perfoliatum* breeding have little genetic diversity from (often) unknown geographic origins (Wever et al. 2019). To date, most *S. integrifolium* breeding progress has been made from accessions that originated within a small geographic area (Kansas, USA) (Vilela et al. 2018), with a general lack of genomic resources (Van Tassel et al. 2017). The extent to which breeding populations are representative of species-wide diversity has not been genetically evaluated. However, if populations that produced *S. integrifolium* breeding accessions are not representative of species-wide diversity, greater improvements may be achieved by incorporating accessions with unique diversity similar to the practice of incorporating new, useful genetic variation into crop breeding programs from their wild relatives (McCouch et al. 2013). Additionally, the choice of locations for field trials may be informed by a better understanding of the relationship between natural genetic variation and geography.

Presently, *Silphium* genomic resources are lacking, in part, because of the difficulty of assembling its large, repetitive genome. All *Silphium* species are diploid (2n = 14) with genome sizes that are nearly triple that of their close relative sunflower (*Helianthus annuus*) (Bai et al. 2012). We utilize transcriptome resequencing of *S. integrifolium* to characterize and quantify functional genetic variation throughout the species range. We assemble a reference transcriptome of a diverse set of tissues from a single individual and use the reference to identify variants from a panel of resequenced young leaf transcriptomes.

Here, we characterize genetic variation among samples of *S. integrifolium*. We use this variation to investigate the distribution of neutral genetic variation across the species range while maintaining an interest in identifying a possible region of geographic origin of the species. In addition to uncovering genome-wide population structure, we hypothesize that populations of this widespread species are locally adapted to their unique environments and thus, quantifying the relationship between genetic variation and geography is important. Therefore, we next sought to identify genes whose patterns of genetic variation have been shaped by natural selection. Such environmentally associated variants highlight traits experiencing geographically variable selection in this species, and could identify important variation to include in the breeding program germplasm.

## Methods

### De novo transcriptome sampling, sequencing, and assembly

Five tissues were collected from a single *S. integrifolium* plant from the breeding program of The Land Institute in Salina, Kansas. Root, crown, flower, leaf, and stem tissues were collected and immediately frozen in liquid nitrogen. Flash frozen samples were transported to the University of Minnesota on dry ice where total RNA was extracted using Qiagen RNeasy kits (Germantown, Maryland USA). Pooled RNAseq stranded libraries were constructed by the University of Minnesota Genome Center and sequenced on two lanes of an Illumina (San Diego, California USA) HiSeq instrument (2 x 125 paired-end reads per lane).

Raw sequence reads were trimmed of low quality sites and adaptor sequences using BBDuk (Bushnell 2018) and *de novo* assembled into a reference transcriptome with Trinity (Haas et al. 2013) (version 2.1.1, non default settings: SS_lib_type_type RF min_contig_length 300 min_kmer_cov 2 min_glue 5).

### Assembled transcriptome completeness and annotation

The completeness of the initial Trinity *S. integrifolium* assembly was measured by assessing the number of benchmark universal single-copy orthologs with the program BUSCO v3.0.2 with the *embryophyta_odb9* gene set (Simão et al. 2015). The longest isoform of each transcript was used for BUSCO searches.

The number of nearly full length protein coding transcripts was counted by querying the Swiss UniProt protein database (Boutet et al. 2016) with the full set of transcripts using BLASTx with an e-value cutoff of 1e-20 (Madden 2013). For each unique result from the protein database, the percent of matching sequence length of the best matching transcript was recorded. Transcripts that matched a database entry over more than 80% of its length were classified as nearly full-length transcripts.

Transcript abundance was estimated with the RSEM software (Li and Dewey 2011) after mapping cleaned sequence reads to the Trinity assembly using Bowtie v1.1.2 (Langmead and Salzberg 2012). Gene abundance matrices were built after calculating the number of gene transcripts per million sampled transcripts (TPM) using Trinity scripts. Transcript abundance matrices per gene were built to determine the isoform prevalence within each gene. The median contig length (N50) of the transcriptome assembly was calculated using all transcripts (E100N50) and only transcripts with expression levels above the 90th percentile (E90N50). Although the N50 statistic is widely reported without regard for expression level, it may be an inaccurate representation of transcriptome assembly because of a bias from low expressed transcripts.

Functional annotation of the longest isoform of each contig was performed using the standard Trinotate pipeline (Bryant et al. 2017). Possible protein coding sequences were identified using TransDecoder (Haas and Papanicolaou 2017). Putative proteins longer than 100 amino acid residues were kept. BLASTx searches using transcripts and BLASTp searches using predicted amino acids as queries were performed against the Uniprot protein sequence database (Madden 2013; Boutet et al. 2016).

### Reference transcriptome filtering

The Trinity assembly was further filtered to create a subset of contigs to serve as a genotyping reference for downstream population genetic analyses. We initially retained only contigs that had a significant BLAST result to a plant after using the longest isoform of each contig as a query to the Swiss UniProt database and the NCBI non-redundant nucleotide database with BLASTp and BLASTn searches, respectively. E-value cutoffs of 1e-20 (BLASTp) or 1e-30 (BLASTn) were used. Plant contigs were identified by keeping queries with a best BLAST result belonging to a genus on the NCBI viridiplantae taxonomy list (Sayers et al. 2009). The completeness of the filtered contig set was assessed by again performing a BUSCO analysis of TransDecoder predicted amino acids with the *embryophyta_odb9* gene set. The plant filtered contig set was further reduced to include only contigs that were inferred to be single copy orthologous genes in both *S. integrifolium* and sunflower, *H. annuus* with OrthoFinder (Emms and Kelly 2015). The resulting set of contigs served as a genotyping reference for downstream resequencing of samples.

### Transcriptome resequencing

Fresh young leaf tissue was collected and flash frozen from 68 wild *S. integrifolium* plants from 39 sampling locations and one *S. lacinatum* plant to be used as an outgroup (Appendix S1; see Supplemental Data with this article). We often have a single or few individuals from each sampling location, however sampling locations are often clustered near each other. Self-incompatible species, such as *S. integrifolium* are expected to have less genetically structured populations and thus few individuals from many sites across a species range may underestimate fine-scale structure (Charlesworth 2003). All plants were grown from wild collected seed in a common garden at The Land Institute in Salina, Kansas. Pooled RNAseq stranded libraries were constructed by the University of Minnesota Genome Center and sequenced using an Illumina HiSeq instrument (2 x 125 PE reads per lane).

### Variant calling workflow

Reads from each resequenced leaf transcriptome were trimmed of adaptor sequences using scythe v0.991 (https://github.com/vsbuffalo/scythe) and aligned to the genotyping reference using STAR v2.5.2b (Dobin et al. 2013). Reads were aligned in a two-pass mode to accommodate the possible mapping of isoforms across splice junctions. PCR duplicates were removed from each alignment using Picard Tools (http://broadinstitute.github.io/picard/). Sample genotypes at variant single nucleotide polymorphisms (SNPs) and non-variant sites were called using Freebayes with all samples assigned to a single population (Garrison and Marth 2012). A minimum site depth of 10 reads was required within each sample for a site to be included in genotype calling.

### Population structure and genetic diversity analyses

A principal components analysis (PCA) was performed on *S. integrifolium* genotypes to assess genetic population structure across the species range. SNPs were filtered to remove rare variants (minor allele frequency (MAF) ≤ 0.05) and sites in linkage disequilibrium (LD) (LD threshold = 0.2, missingness = 0%) prior to performing PCA. We performed a Procrustes transformation of sample principal components (PC1 and PC2) coordinates onto their sampling latitude and longitude coordinates to better visualize the relationship between genetic variation structuring and geographic space (Wang et al. 2010).

The relative effect of isolation-by-distance on the observed population structure patterns was estimated using `conStruct` (Bradburd et al. 2018). Similar to other STRUCTURE-like programs (Pritchard et al. 2000), `conStruct` assigns a proportion of ancestry from different genetic groups (K) to individuals. Additionally, `conStruct` can incorporate a spatial element into its model to account for isolation-by-distance. We ran spatial and non-spatial models over a range of underlying genetic groups (K = 1 through K = 6), and found spatial models always had higher predictive accuracy. We ran multiple spatial models (K = 1 through K = 6) to identify the best K value with a scree-plot of predictive accuracy. For each value of K, we ran four independent runs with a 1000 iteration chain length. Five independent runs, each with four MCMC chains of 5000 iterations were then performed with the inferred optimal K value.

Genetic relationships among *S. integrifolium* samples were inferred using `SVD Quartets` (Chifman and Kubatko 2014). `SVD Quartets` estimates an unrooted phylogeny describing the relationships between sets of four taxa. The correct phylogeny topology under a coalescent process is inferred by singular value decomposition of the genetic distance matrix containing the four taxa. Multiple quartet phylogenies are then assembled into a single phylogeny that contains all taxa. A matrix of biallelic SNPs was used with exhaustive quartet sampling, 100 bootstrap replicates and the Quartet MaxCut tree assembly method to produce an unrooted phylogeny within `Paup` (Swofford 2000). A single individual from Illinois was removed from the SNP data matrix prior to running `SVD Quartets` because we observed evidence that it was of unusually high intraspecific admixed ancestry.

Results from PCA, `conStruct`, and `SVD Quartets` indicated that the major axes of genetic variation separate *S. integrifolium* samples into three clusters that broadly correspond to geographic regions. Hereafter we refer to these broad groupings as the eastern (Mississippi, Arkansas, Missouri, Wisconsin, and Illinois plants), southern (Texas and Oklahoma plants), and western (Kansas and Nebraska plants) regions. When describing finer scale comparisons within regions we identify groups of plants by their American state of sampling origin. These groups were not *a priori* assumed to be populations in the traditional population genetic sense.

The level of support for region-level relationships among groups within *S. integrifolium* across markers was assessed using `TWISST` (Martin and Van Belleghem 2017). `TWISST` prunes a given phylogeny down to one tip from each user-defined taxon group and reports the topology of the pruned tree. The process is repeated and support for each topology is recorded as the number of observations. A maximum likelihood phylogeny was inferred for each contig using RAxML v8.0 (Stamatakis 2014) and each tip was assigned to a regional taxon inferred from the above clustering methods. The pruned topology most often observed for each contig’s reconstructed phylogeny was recorded.

Genetic differentiation among sets of *S. integrifolium* plants grouped by their sampling state of origin was measured by calculating Wright’s F_ST_ (Weir and Cockerham 1984) between all pairwise combinations using a stringently filtered marker dataset with the R package `StAMPP` (Pembleton et al. 2013). Retained SNP markers were biallelic in *S. integrifolium*, had a Phred quality score ≥ 20, had an allele balance within heterozygotes between 0.25 and 0.75 or within homozygotes less than 0.05, had average sequencing depth greater than 10 and less than 200, and MAF ≥ 0.05. One hundred bootstrap replicates were performed across loci to calculate 95% confidence intervals around F_ST_ estimates and to check whether our grouping of plants by sampling state of origin at least reasonably approximates biological populations.

Average numbers of nucleotide differences per base pair at four-fold degenerate sites were calculated between all samples to measure genetic diversity (Nei 1987). We organized the diversity measurements into within (*π*) and between (*D_xy_*) sets of plants grouped by their sampling state of origin. Variation was calculated across contig bootstrap replicates to calculate 95% confidence intervals around mean values. Tajima’s D was calculated within each group of plants on every contig using `vcftools` (Danecek et al. 2011). The level of partitioning of genetic variance was measured using an AMOVA performed on genetic distances (Nei 1972) between individuals with states and regions as hierarchical levels of stratification with the R packages `StAMPP` and `pegas` (Pembleton et al. 2013; Paradis 2010). Statistical significance of variance partitioning among stratification levels was assessed with 10000 random permutations.

We found that samples from Illinois exhibited patterns of nucleotide variation, Tajima’s D, and inferred ancestry that were unexpected within the larger contextual patterns of each metric across the species landscape. We performed the three population *f*_3_ admixture tests in `TreeMix` to test the hypothesis that the Illinois population is the result of past admixture between Wisconsin and a third population instead of the null expectation that the relationships among the sampled populations are the product of a tree-like branching process (Reich et al. 2009; Pickrell and Pritchard 2012). Each test took on the form *f*_3_{IL; WI, X}, where evidence that IL is the result of admixture between WI and X is given by a significantly negative test statistic. For all identities of X, Z-scores and standard errors of estimates were reported from jackknifing 50 SNP blocks. We subsequently removed all Illinois samples from our demographic inference modeling so as to not confuse historical processes with anthropogenic movement.

Data from the three-dimensional joint site frequency spectra was used to infer general branching patterns of *S. integrifolium* regional taxa using `Moments` (Jouganous et al. 2017). For three-dimensional spectra, the tested branching scenarios were those that would result in the tree topologies (East, (South, West)), (South, (East, West)), and (West, (South, East)). For each branching pattern scenario, we fit demographic models with no migration, symmetric migration between each taxa pairs, and asymmetric migration rates. The fit of each model was assessed across 100 replicates using different starting parameters selected from bounded limits decided by trial and error. The likelihood of each model was optimized using Powell’s method and the quality of each model’s fit to the data was assessed with AIC scores. After the best-fitting model was selected for each scenario, uncertainty estimates around each parameter were calculated using the Godambe Information Matrix with 100 bootstrapped spectra (Jouganous et al. 2017).

### Scans for selection

To identify loci possibly under divergent selection, we used BayeScan (Foll and Gaggiotti 2008; Foll 2012) to find statistical outlier SNPs that significantly deviate from neutral expectations. We calculated global F_ST_ scans across sites between regions using the same marker set used to perform an AMOVA. Default BayeScan model parameters were used (burn-in length = 50,000, thinning interval = 10, sample size = 5,000, number of pilot runs = 20, pilot run length = 5,000). Annotation of resulting outlier loci was done using BLASTx, BLASTp and eggNOG searches using Trinotate scripts. We then sought to identify environmental variables that may be associated with the selective pressures acting on each outlier protein. We identified environmental variables that had significant associations with genetic markers within each protein after accounting for the coancestry of samples using BayEnv2 (Günther and Coop 2013). Environmental variables from the geographic midpoint of each population were extracted from the ClimateNA dataset (Wang et al. 2016). Brief descriptions of the climatic variables used are provided in Appendix S3. For each BayeScan identified protein, the Bayes factor, absolute Spearman’s rank, and absolute Pearson correlation coefficients with the largest quantile were used to identify putative environmental variables driving local adaptation.

## Results

### De novo transcriptome and assembly

Trinity assembled 541,120,911 paired-end sequencing reads into 280,573 transcripts and 166,458 contigs, totaling over 246.9 megabases of assembled nucleotides. The median contig length of all transcripts (E100N50) was 1,159bp. The median contig length of transcripts within the top 90th percentile of expression (E90N50) was 2,014bp. The filtered assembly with the longest isoform of each contig totaled 123.5 megabases. A BUSCO analysis of the longest isoform of each assembled contig found that 80.7% of the 1,440 genes in the *embryophyta_odb9* dataset were found in complete form (71.3% as single copy and 9.4% as duplicated). BLASTx searches of the Swiss UniProt database identified 13,143 unique protein results. Of those, 8,418 (64.0%) were found as full-length or nearly full-length transcripts in the full Trinity transcriptome assembly.

### Reference transcriptome filtering

51,238 contigs remained after filtering the full Trinity for contigs with a BLASTp or BLASTn result from genera on the NCBI viridiplantae taxonomy list. Results of a BUSCO analysis of this filtered contig set were similar to the full assembly analyses (80.7% of 1,440 BUSCO genes found complete, of which, 71.4% were single copy and 9.3% were duplicated). The plant-filtered contig set was further reduced to 10,575 contigs that contained orthologous genes that were inferred to be single copy in *Silphium* and sunflower, *H. annuus*. This final contig set was used as a reference to align resequencing reads and to call variants.

### Transcriptome resequencing and variant calling

Leaf transcriptome resequencing data mapped to 7,718 of the 10,575 contigs in the genotyping reference set. Genotyping was restricted to contigs with mapped reads and covered 12,410,795 total sites (852,148 SNPs) across all samples. One *S. integrifolium* sample from Missouri was removed from downstream analyses because of low average sequencing depth (< 10) and a high rate of marker missingness (> 60%).

### Population structure and genetic diversity analysis

PC1 and PC2 explained 13.79% of the variation in our *S. integrifolium* samples. Genotypes formed three distinct groups on the first two PC axes (Fig. 1B). The groups broadly consisted of plants from the eastern (Arkansas, Illinois, Mississippi, Missouri, and Wisconsin), southern (Oklahoma and Texas), and western (Kansas and Nebraska) regions of the species range. Eastern and western regions were separated along the first PC axis. The southern region was separated from eastern and western regions along the second PC axis.

**Figure 1:**
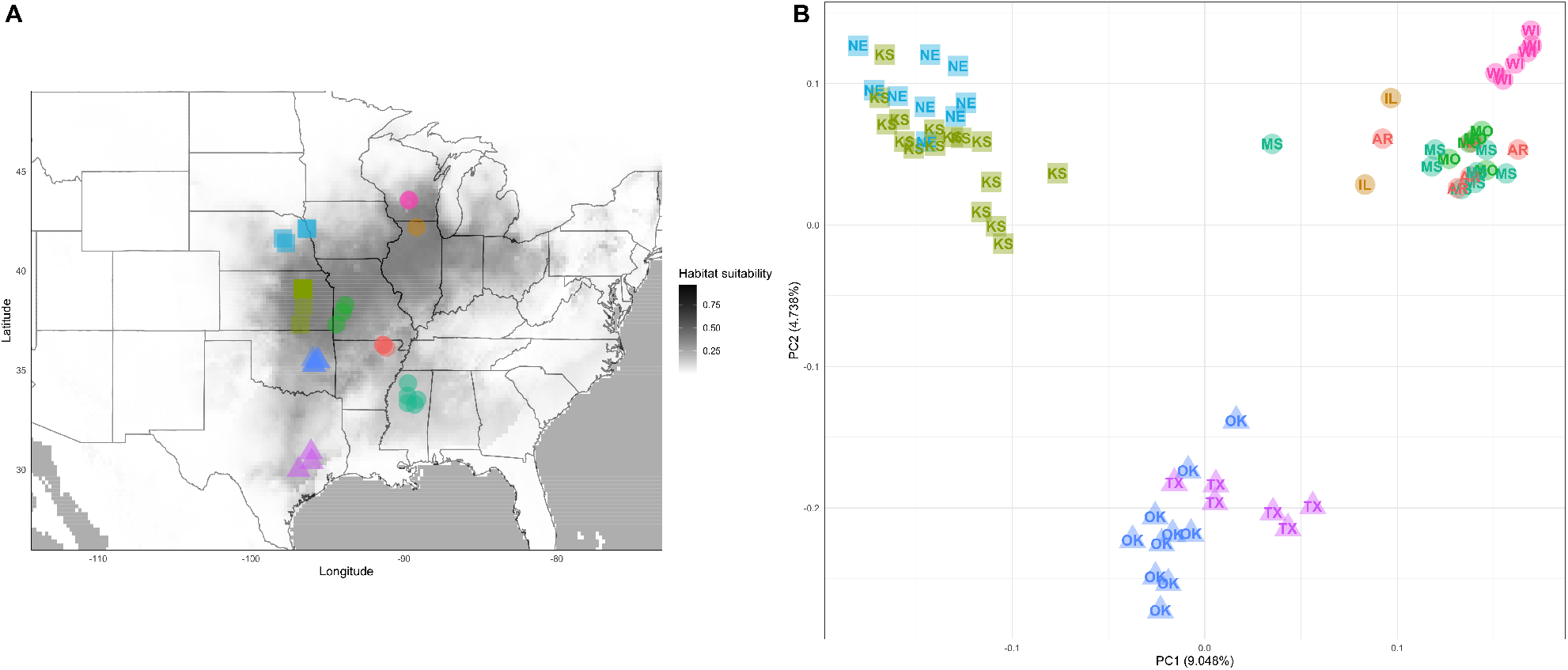
**(A)**: Suitable *S. integrifolium* range estimate and current study sampling locations. Range estimate is shaded by MaxEnt estimated habitat suitability (see Appendix S8 for method details). Sampling localities are colored according to their American state. **(B)**: Results from a principal components analysis of *S. integrifolium* genotypes. The two major axes of variation distinguish samples into three clusters that correspond to regional (eastern, southern, and western) geography. Coloring on principal components biplot corresponds to same states in panel A.

Our `conStruct` analyses that incorporated isolation-by-distance into population structure (spatial models) always performed better than non-spatially aware models. Across ten-fold cross validation runs for each value of K the improvement of spatial over non-spatial models ranged from 291.79 to 54.52 log-likelihood units. Results from spatially aware runs with different numbers of K indicated that relatively little predictive power was gained after increasing the number of genetic layers past K = 3. The patterns of assigned ancestry across all samples supported the PCA pattern of subdivision. Within the eastern region, samples were overwhelmingly comprised of ancestry from a single dominant genetic layer. Samples from the southern and western regions are inferred to be best described as a mixture of ancestrty, with both regions being made up of a different dominant ancestry layer (Figs. 2A, 2B). In western samples, the second largest ancestry layer is the dominant layer in southern samples. The reciprocal is true in most southern samples.

**Figure 2:**
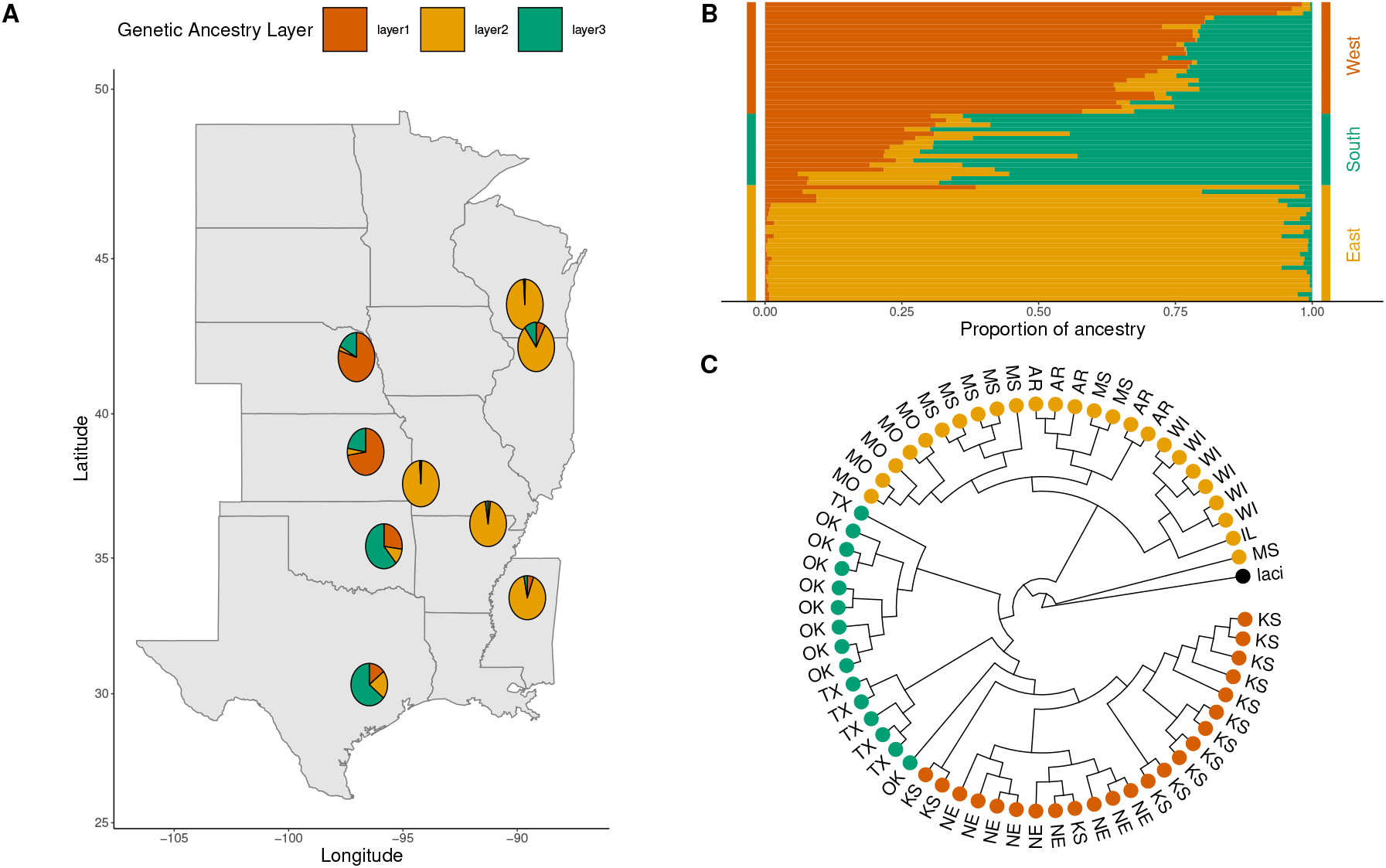
*Silphium integrifolium* samples comprise three genetic groups. **A - B**: Inferred ancestry results of *Silphium integrifolium* samples from a `conStruct` model that includes spatial information and K = 3 (Bradburd et al. 2018). **(A)**: Average amounts of inferred ancestry from three genetic layers for each population are displayed as pie charts. **(B)**: Proportion of inferred ancestry of each genetic layer for all sampled plants. **(C)**`SVD Quartets` reconstruction of phylogenetic relationships between *S. integrifolium* samples, rooted on single *S. laciniatum* sample. Bootstrap node support values are not shown, however every node has 100% bootstrap support.

Reconstruction of the phylogenetic relationships of samples using `SVD Quartets` resulted in strong geographic clustering with 100% bootstrap support for all nodes in the phylogeny (Fig. 2C). All but one sample from the eastern region formed a large monophyletic group, with most samples from within a population forming smaller monophyletic groups. Mississippi samples exhibit a polyphyletic pattern throughout the larger eastern clade, with various samples being found clustered with Missouri or Arkansas samples. A single Mississippi sample was reconstructed to be outside of a larger clade containing all other *S. integrifolium* samples. Southern samples are paraphyletic with most Oklahoma samples forming a monophyletic clade that is sister to a single Texas sample. Most Texas samples form a monophyletic clade that is sister to a larger clade that contains a single Oklahoma sample and a larger monophyletic clade of all western samples (Fig. 2).

To investigate the level of support for the general `SVD Quartets` phylogeny across markers, we performed a `TWISST` analysis. Each maximum likelihood phylogeny was rooted with *S. laciniatum* as the outgroup and the region-level topology of *S. integrifolium* taxa phylogeny was recorded. For each taxa in a tree the identity of its sister tip was recorded. If the nearest sister was a node, a descendant of the sister node was randomly sampled and given a weight proportionate to the number of descendants of the sister node. Confidence around these measurements was estimated with 25,000 bootstrap across contig phylogeny topologies. Similar to the `SVD Quartets` phylogeny topology, southern and western regions were sister to each more often than either was sister to the eastern region. The eastern region was found to be sister to western and southern regions an approximate equal number of times across markers (Appendix S7). For each focal taxon’s set of sister taxa, a chi-square test was performed assuming a star phylogeny i.e. equal probabilities of observing each sister taxon. Equal probabilities of observing each sister taxon was rejected for the western and southern region focal taxa (*p* < 0.05 bonferroni corrected for multiple comparisons).

Across all comparisons, pairwise F_ST_ ranged from 0.0340 to 0.2204. Differentiation patterns between regions supported the structure inferred from PCA. As a general pattern, differentiation within regions was approximately F_ST_ ≤ 0.10 regardless of geographic distance separating samples (Fig. 3). The rates of decay of the covariance between allele frequency and geographic distance within each genetic layer (`conStruct`’s *α*D parameter, K = 3) ranged from 0.001 to 0.005 suggesting little genetic differentiation above what is expected under isolation-by-distance within genetic layers. Differentiation between regions was generally higher than within region comparisons with increased differentiation observed between geographically distant samples (Fig. 3), however the relationship between geographic distance and differentiation is not significant within each type of comparison (within region and between regions). The three highest within-region comparisons all involved the suspect IL locality, likely reflecting the recent admixture in this sample. The highest levels of differentiation were observed between eastern and western regions, and between eastern and southern regions. The greatest allele frequency divergence observed was between Nebraska (western region) and Illinois (eastern region) populations (F_ST_ = 0.2204). Our AMOVA results showed that genetic variation is strongly partitioned both among regions and among states of sampling within regions (Table 1).

**Figure 3:**
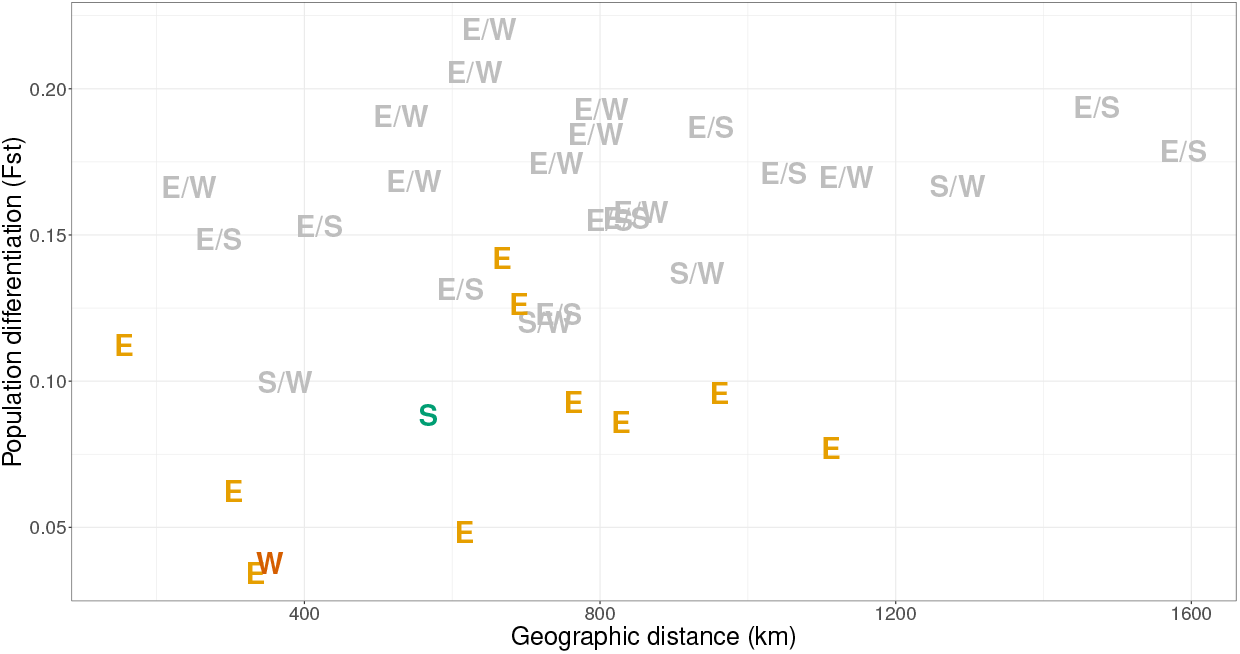
Relationships between geographic distance and genetic differentiation. Geographic distance was measured as Euclidean distance between the sample midpoints of sets of samples grouped by sampling state of origin. Differences in color reference within and between region comparisons; colored letters (E, S, W) represent within region comparisons and gray letter pairs (E/S, E/W, S/W) represent between region comparisons. There is no significant effect of geographic distance on genetic differentiation within each type of comparison (within and between regions, *p* > 0.05.).

**Table 1:** Analysis of molecular variance (AMOVA) based on hierarchical grouping stratification among regions, among populations within regions, and within populations. ∗∗ = *p* ≤ 0.01, ∗ ∗ ∗ = *p* ≤ 0.0001

| Sources of variation | df | SS | Variance components | Percent of variance |
| --- | --- | --- | --- | --- |
| Regions | 2 | 0.44905 | 0.00796** | 29.4869% |
| Populations | 6 | 0.23872 | 0.00375*** | 13.9031% |
| Within Populations | 58 | 0.88744 | 0.01530 | 56.610% |
| Total | 66 | 1.57522 | 0.027028 | 100.00% |

The average numbers of pairwise nucleotide differences per base pair at four-fold degenerate sites (*π*) between samples within regions were calculated to estimate genetic diversity (Fig. 4B, Appendix S2). The eastern region had the highest mean genetic diversity of any group (*π* = 0.01013, 95% C.I. 0.00986 - 0.01041). The southern region had slightly lower genetic diversity (*π* = 0.00982, 95% C.I. 0.00952 - 0.01013). The western region had lowest genetic diversity (*π* = 0.00894, 95% C.I. 0.00865 - 0.00923).

**Figure 4:**
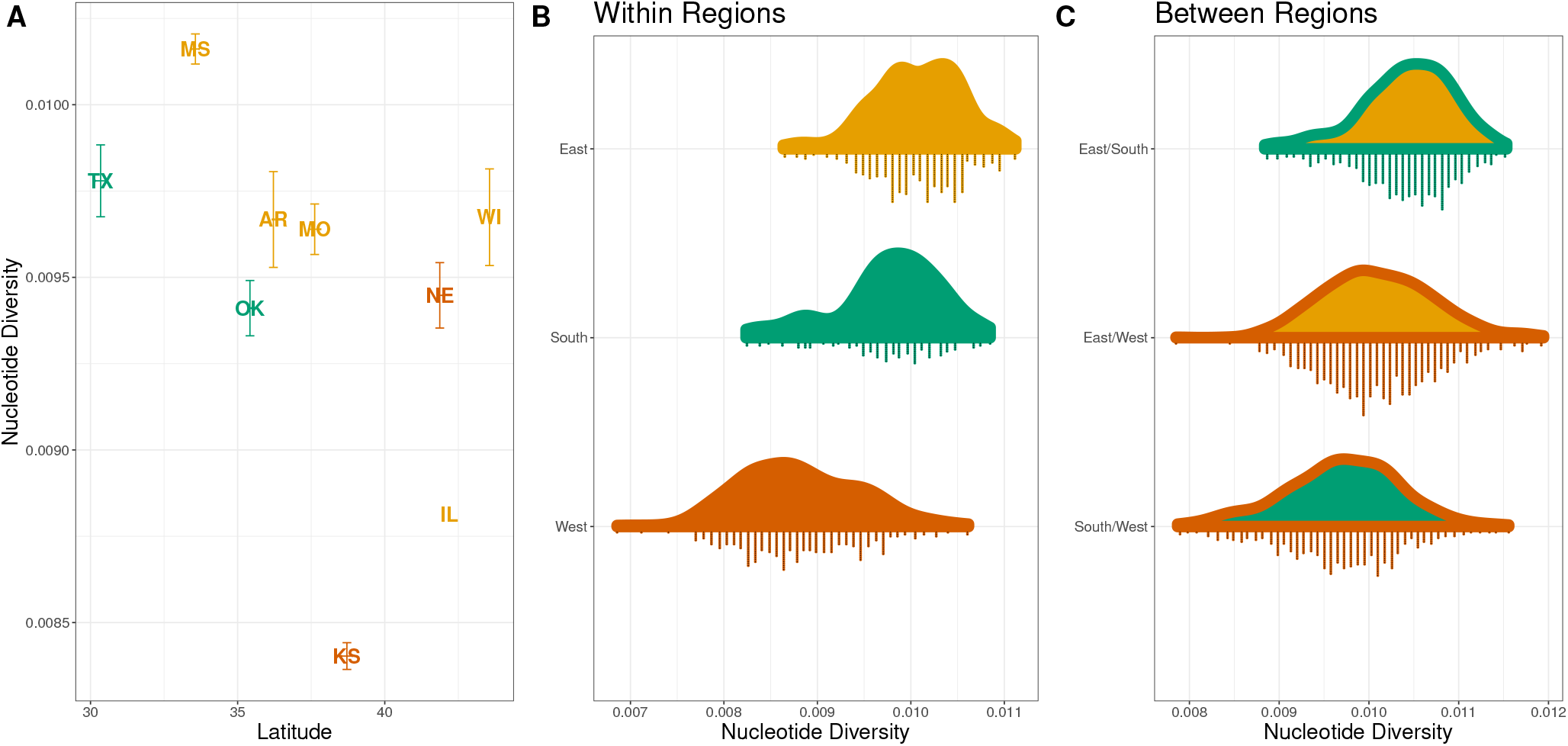
Distribution of nucleotide diversity at four fold degenerate sites. **(A)** Mean nucleotide diversity between samples grouped by sampling state of origin. For each state, latitude is calculated as the latitude midpoint of sampled plants. As a general pattern, diversity decreases with latitude. **(B)** Pairwise nucleotide diversity raincloud plots among samples within regions (*π*) and **(C)** among samples between geographic regions (*D_xy_*). Each raincloud plot displays the nucleotide diversity density function above the observed raw data points. The outline and fill colors of the density distributions correspond to the regions being compared.

Within the eastern and southern regions, samples from states with lower latitudes had higher *π*, consistent with those regions possibly serving as founder populations for northward expansion after glaciation (Fig. 4A, Appendix S2). Mississippi and Arkansas contained more genetic diversity than other eastern region state groupings (*π* = 0.01018, 95% C.I. 0.00987 - 0.01051 and *π* = 0.00972, 95% C.I. 0.00935 - 0.01008, respectively). Within the southern region, genetic diversity in Texas was greater than Oklahoma (*π* = 0.0098, 95% C.I. 0.00944 - 0.01017 and *π* = 0.00949, 95% C.I. 0.00918 - 0.0098, respectively). The reduction in genetic diversity found in Kansas and Nebraska (*π* = 0.00846, 95% C.I. 0.00815 - 0.00878 and *π* 0.00948, 95% 0.00918 - 0.00978, respectively)) is suggestive of a past bottleneck of *S. integrifolium* into the western region.

Patterns of nucleotide divergence between populations (*D_xy_*) support the regional structure grouping of plants found through PCA and `conStruct`. The largest *D_xy_* observed, on a regional scale, was between eastern and southern regions (*D_xy_* = 0.0105, 95% C.I. 0.01021 - 0.0108), only slightly exceeding *π* in the diverse eastern region. The largest levels of divergence were found between Mississippi and Texas (*D_xy_* = 0.01069, 95% C.I. 0.01036 - 0.01102) followed by Texas and Wisconsin (*D_xy_* = 0.01068, 95% C.I. 0.01035 - 0.01102), and Mississippi and Nebraska (*D_xy_* = 0.01067, 95% C.I. 0.01036 - 0.01098). Plants from the western region had the smallest divergence from southern region plants, suggesting that the western samples are a result of a founder event originating from the southern region (Figs. 3, 4C, Appendix S2).

Tajima’s D, calculated within each state across contigs, was consistently negative within samples from eight of nine states (Fig. 6), suggesting population growth across the range. We note that this runs counter to the known recent contraction of prairie species due to habitat loss, and likely reflects population growth since the glacial retreat. A recent selective sweep can also cause negative Tajima’s D, however the genomic signature will be contained to genetic markers linked to the target of the sweep. Yet another alternative is this negative Tajima’s D reflects population substructure within states, however the distribution of pairwise *π* within states argues against this explanation. The genome wide pattern of negative Tajima’s D we observe is thus interpreted to be due to population expansion, a process that will affect markers across the genome. The western region, (Kansas and Nebraska), Oklahoma, and Mississippi samples had the most negative Tajima’s D values. Illinois was the only grouping that had an average Tajima’s D value greater than zero (D = 0.201), likely reflecting the excess of intermediate frequency variants induced by rampant recent admixture.

The relatively high genetic variation (*π*), elevated Tajima’s D estimates, and amounts of mixed ancestry inferred by `construct` within the IL population led us to test for a history of admixture using the *f*_3_ test for treeness (Reich et al. 2009). We assumed that the IL and WI samples have recently diverged from each other via a common ancestral population and thus tested the null hypothesis that a tree-like branching pattern would best describe the process that formed any trio of populations containing IL and WI in the form *f*_3_{IL; WI, X}. We found significantly negative Z-scores for the *f*_3_ test when the third population in the trio was any western or southern grouping (KS, NE, OK, or TX), but not any other eastern grouping (AR, MO, or MS) (Table 2). Overall, this pattern suggests that IL is the product of admixture between the general eastern region lineage and the western or southern regions lineages or the common ancestor of the two.

**Table 2:** Results from *f*_3_ tests for introgression to test the hypothesis that the Illinois population is admixed. Significantly negative test statistics indicate that IL population has history of a non tree-like branching process *i.e*. past admixture between populations P1 and P2.

| Focal | P1 | P2 | $f_3$ | std. error | Z-score | p value |
| --- | --- | --- | --- | --- | --- | --- |
| IL | WI | MS | -0.0012 | 0.0010 | -1.1329 | 1.286e-01 |
| IL | WI | AR | -0.0004 | 0.0011 | -0.3531 | 3.620e-01 |
| IL | WI | MO | -0.0004 | 0.0010 | -0.3724 | 3.548e-01 |
| IL | WI | TX | -0.0031 | 0.0011 | -2.7611 | 2.881e-03 |
| IL | WI | OK | -0.0042 | 0.0012 | -3.5755 | 1.748e-04 |
| IL | WI | KS | -0.0045 | 0.0010 | -4.3112 | 8.118e-06 |
| IL | WI | NE | -0.0044 | 0.0011 | -4.1234 | 1.867e-05 |

Data from the joint site frequency spectra was used to model the demographic history of *S. integrifolium* under three general scenarios. The scenarios differed from each other by the identities of the two regional taxa assumed to be sister to each other *e.g*. (East, (West, South)), (South, (East, West)), and (West, (East, West)). Under each model, an ancestral population was forced to split into two lineages, with one lineage later experiencing a second split to produce three total taxa. The proportions of a lineage that went into a split at a branching event, timing of branching events, migration rates between all lineages, and population sizes were estimated for each model. Migration rates differed in each model to either be nonexistent, symmetric, or asymmetric.

Under all branching scenarios considered, models that included migration were preferred over no-migration models. Models with migration rates constrained to be symmetric were preferred over asymmetric migration models. The three different branching scenarios considered for three-dimensional spectra each resulted in similar maximum likelihoods and AIC scores (Fig. 5, Table 3). Overall, the best fitting model produced the (East, (South, West)) topology with substantial population expansion in the South and West with migration throughout all populations. This model estimates that approximately 40% of the ancestral population split ≈ 3.7*N_e_* generations ago to form the common ancestor of South and West regions. This lineage then approximately doubled in size before splitting again ≈ 1.6*N_e_* generations ago to create the South and West regions. At the most recent branching event, the majority of the population (83.7%) split to form the South region, which has since decreased by nearly half. The West region’s population size has increased nearly fourfold since the most recent split. The population of the East region has been stable in size since its initial split from the common ancetor. Estimated migration rates are highest between East-South and South-West regions, however confidence intervals around migration rates are wide. Parameter estimates under all models are reported in Table 3.

**Figure 5:**
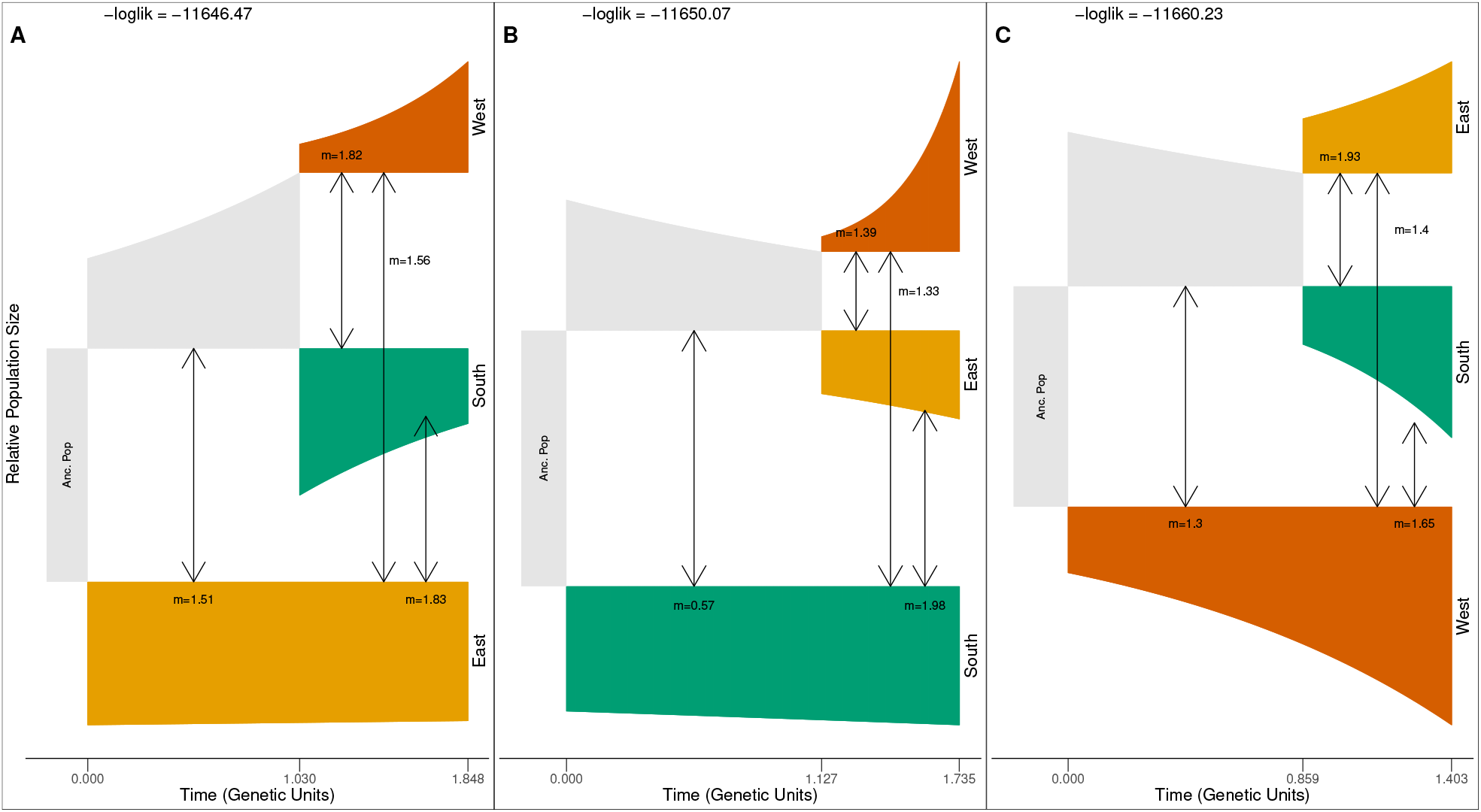
Summarized results of best fitting demographic inference model (symmetric migration rates) for each lineage branching scenario, arranged left to right from highest to lowest log-likelihood. For each plot, sampled extant lineages are colored and ancestral lineages are gray. Each simulation evolves from an initial ancestral population (left most gray box) and undergoes two splitting events to produce three extant lineages (right edge of each plot). The log-likelihood is reported above each plot. Height of each lineage segment is equal to population size relative to ancestral population (Anc. Pop) size. Migration rate estimates are reported near arrows in each figure. Estimated model parameters with standard deviations and model fit statistics are listed in Table 3. Time and migration rates are in genetic units (2*N_e_*).

**Table 3:**
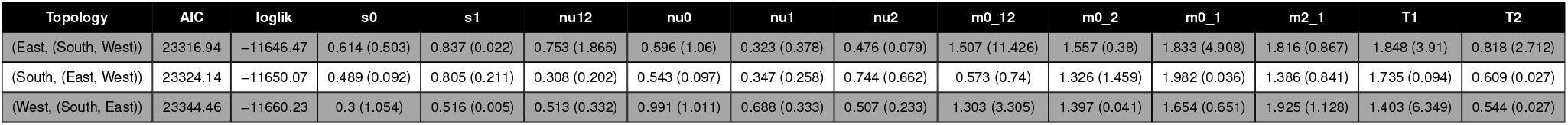
Demographic inference model parameter estimates with standard deviations. For each scenario topology *e.g*. (0,(1,2)), *s* parameters are proportion of lineage that split into population (*s*0 = proportion of ancestral population that split into population 0), *nu* parameters are population sizes, *m* parameters are migration rates between listed populations, *T*1 is time since initial split, *T*2 is time since second split.

### Scans for selection

We performed a transcriptome wide scan for divergence outliers on a set of 41,159 high quality markers with BayeScan. We detected 47 SNPs that were F_ST_ outliers while using a maximum false discovery rate of 0.001. The outlier SNPs were spread across 40 contigs, 34 of which we were able to gather annotation information from BLAST and eggNOG database searches. The summarized annotation information for selected proteins are given in Appendices S4, S5, and S6. For each protein identified by BayeScan as containing an outlier SNP, we found the environmental variable that had the highest correlation with any SNP based on Bayes factor, Spearman rank correlations, or Pearson correlation coefficient quantiles using BayEnv2. We speculated that the combined knowledge of the protein function from BLAST and eggNOG searches and associated environmental variables would lead to better understanding past selective pressures *S. integrifolium* experienced while expanding to its current geographic range. Several genes with known functions associated with abiotic stress tolerance show associations with climatic variables. Below, we highlight environmental variables that are associated with genes to further understand biological processes that may have enabled range expansion. For example, Delta(8)-fatty-acid desaturase has a role in tolerating cold and chilling environments (Chen et al. 2012) and was found to be associated with both continentality (temperature difference between warmest and coldest months) and mean annual precipitation variables. Aquaporin TP1 encodes a channel protein involved in water transport that has been shown to influence cell turgor and expansion (Maurel et al. 1993) and was found to be associated with annual heat moisture index, a composite measure of heat and precipitation, and longitude. The E3 ubiquitin ligase SUD1 gene is involved in drought tolerance through its effects on cuticle wax formation and transpiration rate (Lü et al. 2012) and was found to be associated with the mean warmest month temperature and extreme warm temperatures.

## Discussion

### Reference transcriptome as a resource

The development of *Silphium* genomic resources has been hindered because of difficulties imposed by its large genome that is replete with repetitive elements. We sought transcriptome resequencing as a reliable method to obtain a common set of reduced representation genetic markers from this non-model organism. Towards this goal, we built a reference transcriptome of a diverse set of tissues from a single plant and used it as a genotyping reference for the resequencing of a species wide panel of *S. integrifolium* samples. The initial transcriptome assembly contained roughly 250Mb of sequence data assembled into more than 166,000 contigs, some of which are products of sequence contamination and assembly error. An assembly of the longest isoform of each contig contained a majority of nearly full-length transcripts and an almost complete set of universal benchmark genes predicted to be present in all plants. Our final genotyping reference was the product of filtering to include only contigs that contained genes that have been sequenced in other plants and present as single copies in both *Silphium* and its close relative, sunflower. Assuming approximate conservation of the gene number present across plants, the reference transcriptome contains roughly one third of genes in *Silphium* and can presumably serve as a helpful genotyping resource in place of a quality reference genome (Sterck et al. 2007).

### S. integrifolium comprises three genetic groups

We resequenced the young leaf transcriptomes from dozens of *S. integrifolium* plants from localities that cover the species range. Multiple analyses to identify genetic population structure across samples found that *S. integrifolium* comprises three clusters (Figs. 1, 2). Although the genetic clusters are each found within particular geographic regions of the United States, there are no obvious physical barriers that may help maintain cluster integrity by restricting gene flow. Groups of samples that are geographically near (≈200 km) but belong to different regional groupings show greater differentiation, as measured by F_ST_, than the most distant (∼1000 km) pairs from the same regional grouping (Fig. 3). Although we found considerable levels of neutral genetic variation within each population or region, both are lower than levels found in other self-incompatible wild sunflower species (Fig. 4). Using a set of nuclear genes, Liu and Burke (2006) found approximately three times the amount of synonymous variation in wild *H. annuus* (wild populations *π* = 0.0315) than what we observe in *S. integrifolium*. The discrepancy suggests that *S. integrifolium* experienced a much smaller historical effective population size than the annual sunflower, perhaps reflecting *S. integrifolium*’s longer generation time as a long-lived perennial and/or other features of its life history (Andreasen and Baldwin 2001).

Plants from the eastern region form the most genetically variable cluster (Fig. 4B). Within the eastern region there is a general negative correlation between latitude and *π* (Fig. 4A). The Illinois samples were found to have appreciable amounts of admixed ancestry (Figs. 2A, 2B). Results from *f*_3_ tests for introgression having a role in the Illinois population’s past were significant (Table 2). Introgression from the west could potentially have proceeded naturally through unsampled populations, *e.g*. Iowa, or from western *S. integrifolium* seed introduced by humans in prairie restoration projects. Within the eastern region, populations are separated by latitude in phylogenetic analyses (Fig. 2C).

The southern region maintains a breadth of genetic variation that is comparable to the variation of the eastern region (Fig. 4B). There is a negative correlation between latitude and genetic diversity across the two states. All southern region samples were found to be well described by a mix of its own and other ancestry (Figure 2B). The Texas samples are best described as a mixture of ancestry from the South and Eastern regions, and the Oklahoma samples described as a mixture of ancestry from the South and Eastern regions. We note that this description does not necessarily imply recent gene flow or admixture.

The western region contains the least amount of nucleotide variation (Figure 4B). The same negative correlation between latitude and genetic variation is observed in western samples (Figure 4A). Almost all western samples were inferred to be overwhelmingly made up of a single ancestry group (Figure 2B). The reduced levels of polymorphism in the west result in strong phylogenetic clustering that largely groups with geography. Kansas samples are paraphyletic with Nebraska samples forming a monophyletic clade that is embedded within the larger western clade. All western samples form a monophyletic clade that is sister to a sample from the Oklahoma population (Figure 2C).

### Geographic origin of S. integrifolium

It has been claimed that *S. integrifolium* originated from within the historical range of the American tallgrass prairie, *i.e*. the “prairie origin hypothesis” (Settle and Fisher 1972). The close association of the species with the prairie habitat and the inferred young age of the species were cited as evidence for this assertion. More recently, a phylogenetic treatment of the Silphium genus reconstructed *S. integrifolium* to be most closely related to the common prairie cup plant *S. perfoliatum*, and *S. wasiotense* (Clevinger and Panero 2000). Medley (1989) used populations endemic to the dry-mesic, mixed hardwood forests of eastern Kentucky to describe *S. wasiotense* and estimated the species to be closely related to other congeners found on limestone and sandstone soils throughout the unglaciated Cumberland Plateau (*S. brachiatum* and *S. mohrii*). The present study represents the first application of population genetic data to the topic of identifying the geographic origin of *S. integrifolium*. Our results contradict the “prairie origin hypothesis” and suggest that the geographic origin of *S. integrifolium* is in the American southeast.

We hypothesize that *S. integrifolium* has expanded its range from unglaciated regions near present day Mississippi or Texas to cover the American prairie *i.e*. “refugium origin hypothesis”. The evidence for our refugium origin hypothesis is multifold. First, there is a consistent correlation between latitude and diversity. Mississippi plants are the most genetically diverse samples. Texas plants, from the southern region, are also very diverse. However the level of divergence found between the two groups of plants is within the range of variation found within Mississippi, but not Texas. From the sequence divergence data, this pattern suggests that Texas plants are derived from a population somewhat like our Mississippi sample, however more sampling is required to better understand the progenitor taxon in this split. Results from demographic modeling using three-dimensional joint site frequency spectra generally support the refugium origin hypothesis (Figure 5), however confidence intervals around exact parameter estimates are broad (Table 3). We hypothesize that after the split between Mississippi and Texas populations, both eastern and southern lineages expanded northward, reducing genetic variation with northward progression. The expansion of Oklahoma populations into Kansas and Nebraska resulted in a large reduction in genetic diversity. This reduction in variation appears to be most recent and results in western samples still having considerable amounts of the ancestry layer that is dominant in southern samples (Figure 2A, 2B).

The overall negative values of Tajima’s D in all groups, excluding Illinois, suggest a relatively recent population expansion for the entire species. We interpret the patterns of Tajima’s D across contigs to help us identify the most recent population expansion. Western and southern regions have the largest negative values of Tajima’s D, indicating a large excess of rare variants possibly due to a very recent population expansion after bottlenecks. The smaller negative Tajima’s D values found in the other eastern region may be the result of a more modest and/or a more ancient expansion, after which genetic variation has started to accumulate at sites with initially rare variants.

The results of our phylogenetic analyses support the refugium origin hypothesis. We observe a series of nested relationships within the `SVD Quartets` phylogeny (Fig. 2C). Western samples form a monophyletic clade that is nested within a larger clade also containing southern samples. The larger clade containing all southern and western samples is sister to an eastern sample clade. Based on pubescence and number of ray florets of our specimens (unpublished data) we suggest that the eastern clade corresponds to the subspecies *S. integrifolium var. integrifolium* and the other clade corresponds to the subspecies *S. integrifolium var. laeve* (http://www.efloras.org). The high levels of observed variation in Mississippi are consistent with samples being placed throughout the eastern region clade and outside of the larger *S. integrifolium* clade. The results of our `TWISST` analyses suggest that the general topological pattern we observe in our `SVD Quartets` phylogeny has support across the majority of transcriptome markers. The southern and western regions are found to be sister taxa to each other more often than either is to the eastern region taxon. This pattern is consistent with our demographic inference results that western and southern regions have diverged most recently within this species, and that their common ancestor split from an eastern region population (Fig 5). Southern and western regions are found to be sister to the eastern region in equal proportions. This pattern is expected under the refugium origin hypothesis.

The two competing hypotheses are not necessarily mutually exclusive. The prairie origin hypothesis (Settle and Fisher 1970, 1972) coupled with the historical range of the prairie (Kurtz 2013) implicitly suggested to many that the central prairie was the center of *S. integrifolium* diversity. At its largest expanse, the historical prairie included a small amount of habitat in present day Mississippi that was geographically separated from the larger contiguous prairie (Kurtz 2013). We can not rule out that ancestral *S. integrifolium* originated in the small, restricted prairie habitat of Mississippi. However, an origin in the geographic center of the current prairie is not consistent with the genetic variation observed. The best remaining hypotheses, given our observed patterns of genetic variation, would be an origin in the southeastern edge of its current range. A more thorough sampling of Mississippi and Texas *S. integrifolium* and wide geographic sampling of *S. wasiotense*, and *S. perfoliatum* are recommended to further investigate the geographic origin of *S. integrifolium* and interspecific relationships.

### Potentially adaptive candidate genes

Outlier analyses based on BayeScan indicate that several dozens of the genes with the highest average divergence have SNPs that are statistical outliers, falling outside the distribution of most transcriptome markers. Several of these genes show interesting annotations. Together, these candidate loci may have been involved in adaptation to the drier environment with more extreme temperatures experienced during *S. integrifolium*’s hypothesized expansion out of the American southeast.

The SNPs with extremely low false discovery rate (q < 0.001) have patterns of genetic variation highly unlikely to have occurred by chance. The best explanation for these patterns is that they are very tightly linked to genetic variation under selection during local adaptation, and indeed may be in genes important for adaptation. These genes encode proteins involved in transcriptional regulation (Transcriptional corepressor SEUSS), secondary compound metabolism, disease tolerance, and abiotic stress responses (Delta(8)-fatty-acid desaturase, aquaporin TP1), all of which could have been important for adapting to local conditions. In order to better understand the local environment facets that were potential drivers of adaptation, we looked for climatic variables that were correlated with each selected protein across the landscape. The large number of mitochondrial and chloroplast targeted proteins suggests a possible role for cytonuclear interactions. The presence of several genes involved in cytokinin responses may be explained by altered root-shoot ratios being important in developing the extreme drought tolerance for which this species is known. In addition to being candidate genes that may underlie local adaptation, the high divergence at many of these proteins make them interesting to examine further as targets for breeding, particularly in cases where they may alter phenotypes important for domestication. As an example, the variation found in 13S globulin seed storage protein 2 is potentially important because in other species this protein family is a major driver of seed nutritional content.

### Including diversity into breeding program

Perhaps motivated by the prairie origin hypothesis, the overwhelming majority of germplasm involved in *S. integrifolium* breeding programs derive from collections from the western region (Vilela et al. 2018). From those limited collections, significant heritable domestication improvements have been made in the accumulation of above ground biomass and seed yield (Vilela et al. 2018). Our population genetic results suggest that past domestication improvements have been achieved with a limited genetic variation resource. The western region, while containing considerable genetic variation, is depauperate of variation relative to the eastern and southern regions. Evidence for population recovery from a recent genetic bottleneck (e.g., low Tajimas D values, Fig. 6) could imply that potentially useful variation may have been lost during adaptation to this drought prone climate.

**Figure 6:**
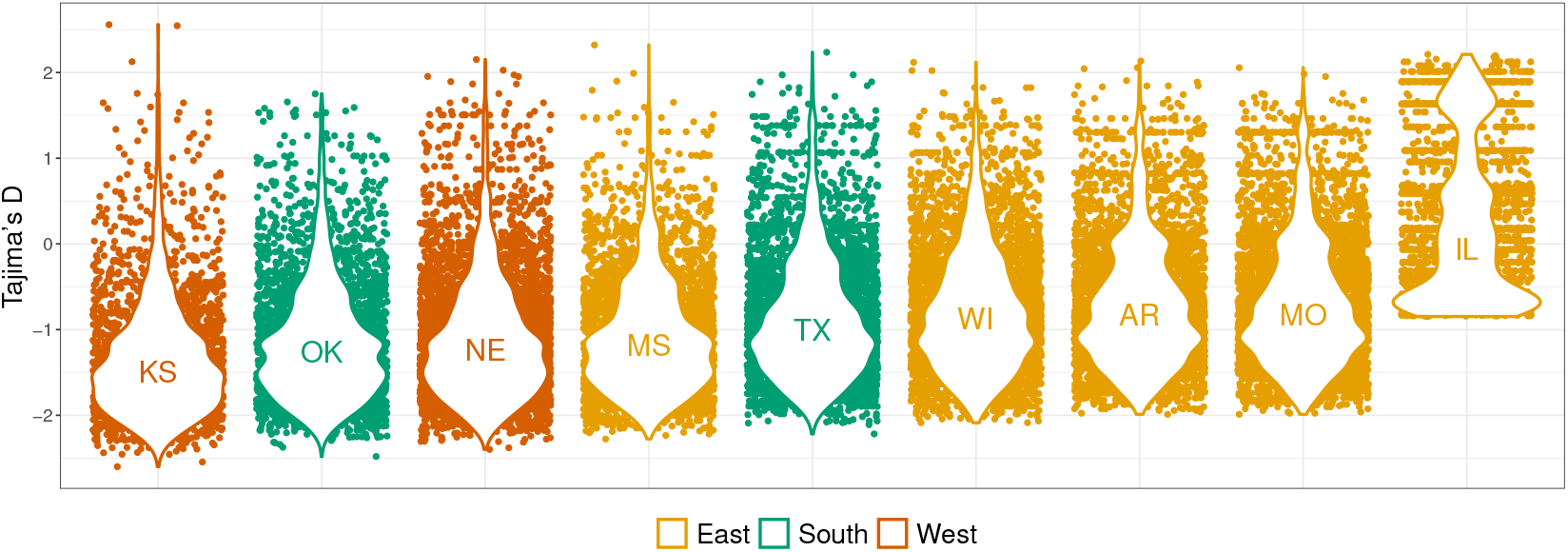
Distribution of Tajima’s D across contigs among grouped plants within each sampling state of origin. Colors correspond to different regional groupings inferred from PCA, `conStruct`, and `SVD Quartets` analysis. Across markers the majority of Tajima’s D estimates are negative, consistent with a recent species-wide range expansion. We hypothesize that the elevated Tajima’s D found in Illinois samples are the result of recent introgression between Illinois and non-Eastern region plants.

The elevated genetic variation across the rest of the species range and the increased divergence between regions is evidence that useful variation that may be beneficial to domestication efforts can be further incorporated into the breeding program. Characterizing the potential primary gene pool of a crop can help breeders identify useful parents in at least two ways. First, demonstrating trait or allele association with climatic, soil or geographic variables could enable breeders to predict which wild populations might produce individuals with extreme values for traits of interest. Recent studies have found that the domestication-targeted traits, seed size and seed oil profile, in wild populations have clear geographic patterns (Reinert et al. 2019). Second, information about the distribution of genetic variation within and between populations and phylogenetic relationships between populations may inform the sampling strategy for a wild germplasm collection campaign designed to capture the most genetic variation in the least number of accessions that need to be subsequently characterized and maintained. The kind of genetic variation thus preserved ex situ may require careful screening (*e.g*. for resistance to a particular pathogen) to be valuable to breeders, or may be held in reserve for future breeders seeking new genetic variation to adapt crops to evolving pests or climate change. This information should also help inform in situ conservation strategies, and in this case it argues for high priority to be placed on conserving rare native prairies in the southern United States (Campbell and Seymour 2012) as these appear to be the center of origin of at least one *Silphium* species and perhaps a similar pattern will be found for other native grassland species.

## Acknowledgements

The authors thank (1) the associate editor and anonymous reviewers for their helpful comments and suggestions that improved the manuscript, (2) John Holmquist, who found and collected many of the wild populations and helped with RNA extractions, (3) other members of the public, botanical gardens, and prairie conservation organizations who have donated wild *Silphium* germplasm to the The Land Institute’s collection over the years, (4) the financial support of the Perennial Agriculture Project and The Land Institute’s donors, (5) Kelsey Peterson, John Hill Price, and Kevin P. Smith for their discussion and advice which improved the manuscript, and (6) NSF grant #1737827 Dimensions US-China to YB.

## Supporting information

Additional Supporting Information may be found online in the supporting information section at the end of the article.

## Appendices

**Appendix S1:**
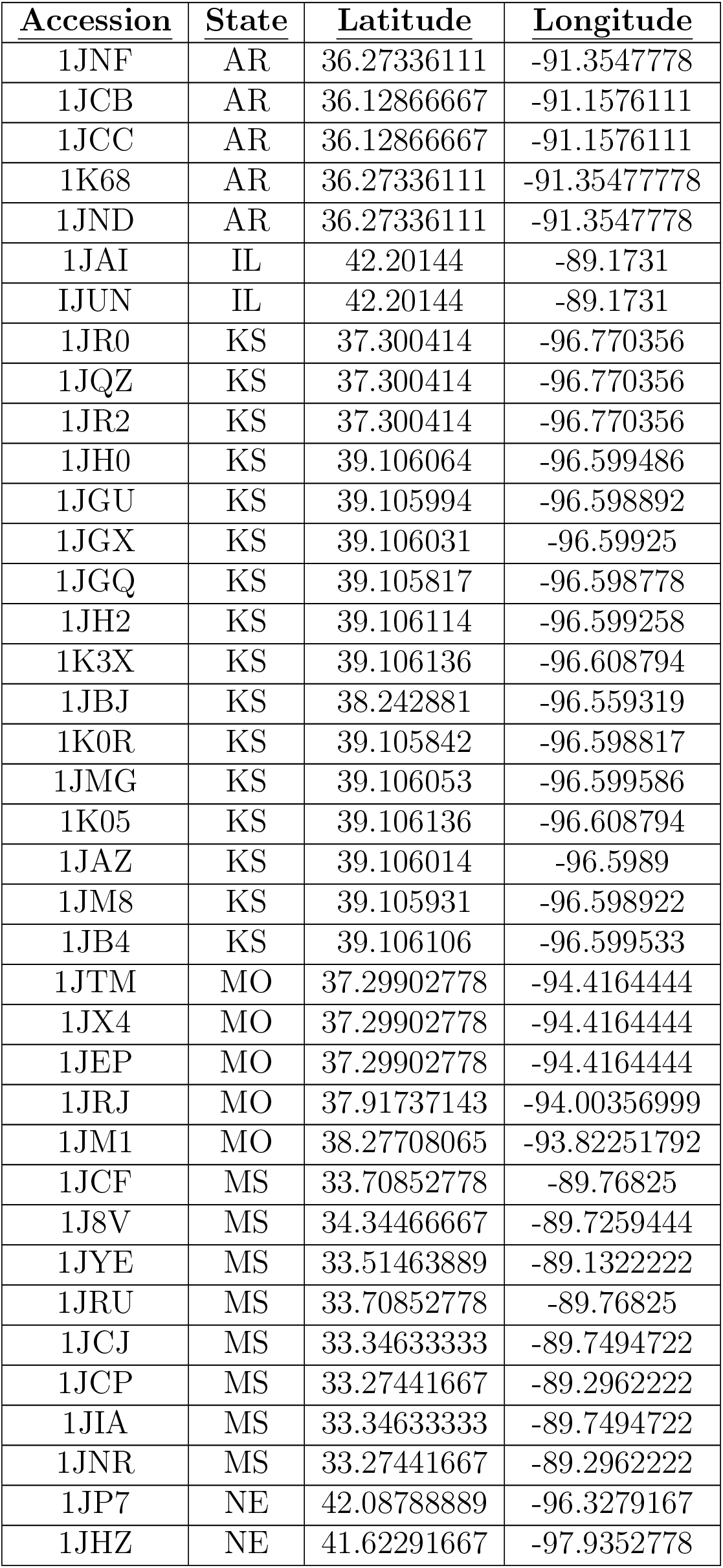
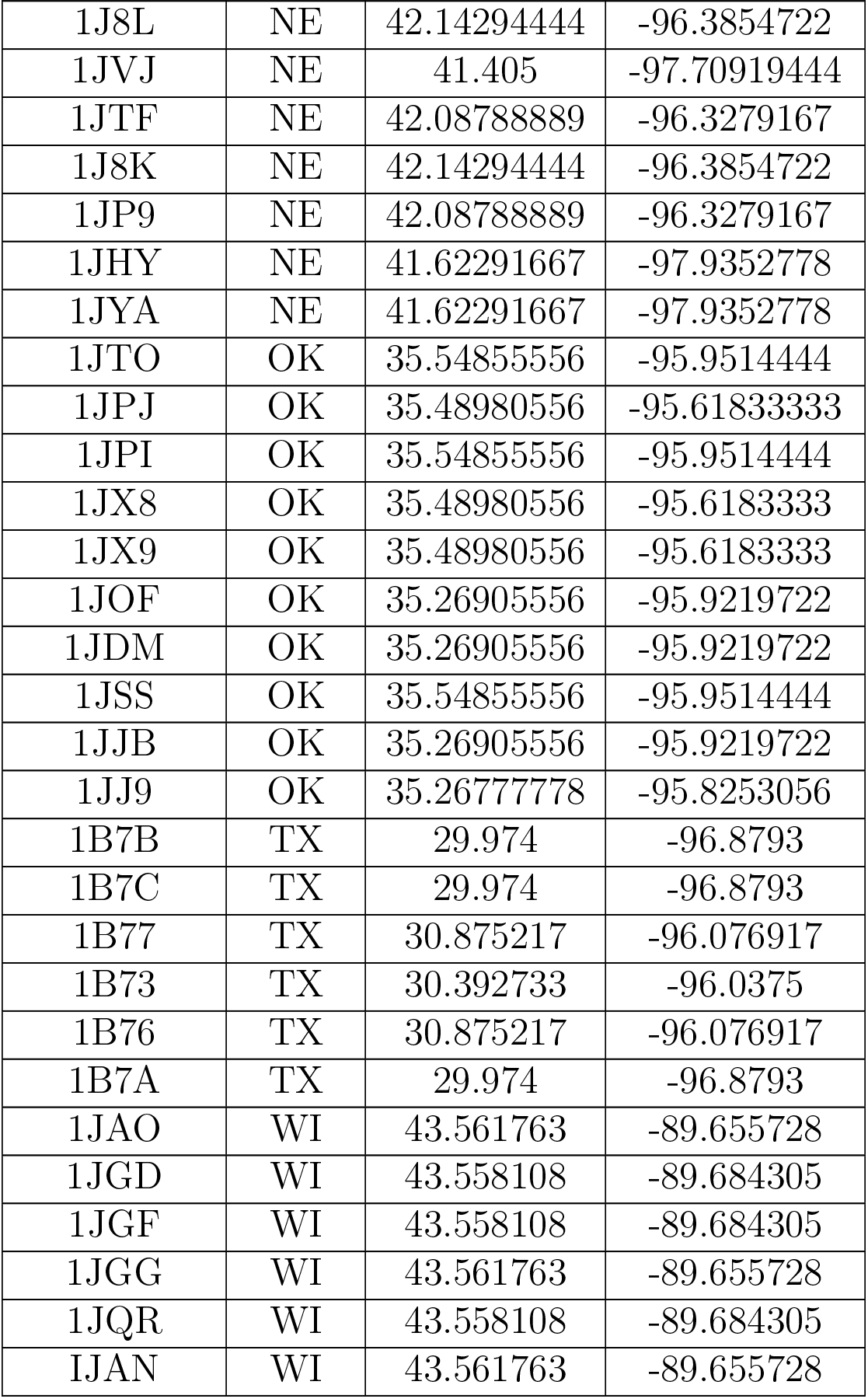
Geographic locality for all *S. integrifolium* samples used in transcriptome resequencing study.

| **Accession** | **State** | **Latitude** | **Longitude** |
| --- | --- | --- | --- |
| 1JNF | AR | 36.27336111 | -91.3547778 |
| 1JCB | AR | 36.12866667 | -91.1576111 |
| 1JCC | AR | 36.12866667 | -91.1576111 |
| 1K68 | AR | 36.27336111 | -91.35477778 |
| 1JND | AR | 36.27336111 | -91.3547778 |
| 1JAI | IL | 42.20144 | -89.1731 |
| 1JUN | IL | 42.20144 | -89.1731 |
| 1JR0 | KS | 37.300414 | -96.770356 |
| 1JQZ | KS | 37.300414 | -96.770356 |
| 1JR2 | KS | 37.300414 | -96.770356 |
| 1JH0 | KS | 39.106064 | -96.599486 |
| 1JGU | KS | 39.105994 | -96.598892 |
| 1JGX | KS | 39.106031 | -96.59925 |
| 1JGQ | KS | 39.105817 | -96.598778 |
| 1JH2 | KS | 39.106114 | -96.599258 |
| 1K3X | KS | 39.106136 | -96.608794 |
| 1JBJ | KS | 38.242881 | -96.559319 |
| 1K0R | KS | 39.105842 | -96.598817 |
| 1JMG | KS | 39.106053 | -96.599586 |
| 1K05 | KS | 39.106136 | -96.608794 |
| 1JAZ | KS | 39.106014 | -96.5989 |
| 1JM8 | KS | 39.105931 | -96.598922 |
| 1JB4 | KS | 39.106106 | -96.599533 |
| 1JTM | MO | 37.29902778 | -94.4164444 |
| 1JX4 | MO | 37.29902778 | -94.4164444 |
| 1JEP | MO | 37.29902778 | -94.4164444 |
| 1JRJ | MO | 37.91737143 | -94.00356999 |
| 1JM1 | MO | 38.27708065 | -93.82251792 |
| 1JCF | MS | 33.70852778 | -89.76825 |
| 1J8V | MS | 34.34466667 | -89.7259444 |
| 1JYE | MS | 33.51463889 | -89.1322222 |
| 1JRU | MS | 33.70852778 | -89.76825 |
| 1JCJ | MS | 33.34633333 | -89.7494722 |
| 1JCP | MS | 33.27441667 | -89.2962222 |
| 1JIA | MS | 33.34633333 | -89.7494722 |
| 1JNR | MS | 33.27441667 | -89.2962222 |
| 1JP7 | NE | 42.08788889 | -96.3279167 |
| 1JHZ | NE | 41.62291667 | -97.9352778 |

| <u>Accession</u> | <u>State</u> | <u>Latitude</u> | <u>Longitude</u> |
| --- | --- | --- | --- |
| 1J8L | NE | 42.14294444 | -96.3854722 |
| 1JVJ | NE | 41.405 | -97.70919444 |
| 1JTF | NE | 42.08788889 | -96.3279167 |
| 1J8K | NE | 42.14294444 | -96.3854722 |
| 1JP9 | NE | 42.08788889 | -96.3279167 |
| 1JHY | NE | 41.62291667 | -97.9352778 |
| 1JYA | NE | 41.62291667 | -97.9352778 |
| 1JTO | OK | 35.54855556 | -95.9514444 |
| 1JPJ | OK | 35.48980556 | -95.61833333 |
| 1JPI | OK | 35.54855556 | -95.9514444 |
| 1JX8 | OK | 35.48980556 | -95.6183333 |
| 1JX9 | OK | 35.48980556 | -95.6183333 |
| 1JOF | OK | 35.26905556 | -95.9219722 |
| 1JDM | OK | 35.26905556 | -95.9219722 |
| 1JSS | OK | 35.54855556 | -95.9514444 |
| 1JJB | OK | 35.26905556 | -95.9219722 |
| 1JJ9 | OK | 35.26777778 | -95.8253056 |
| 1B7B | TX | 29.974 | -96.8793 |
| 1B7C | TX | 29.974 | -96.8793 |
| 1B77 | TX | 30.875217 | -96.076917 |
| 1B73 | TX | 30.392733 | -96.0375 |
| 1B76 | TX | 30.875217 | -96.076917 |
| 1B7A | TX | 29.974 | -96.8793 |
| 1JAO | WI | 43.561763 | -89.655728 |
| 1JGD | WI | 43.558108 | -89.684305 |
| 1JGF | WI | 43.558108 | -89.684305 |
| 1JGG | WI | 43.561763 | -89.655728 |
| 1JQR | WI | 43.558108 | -89.684305 |
| IJAN | WI | 43.561763 | -89.655728 |

**Appendix S2:**
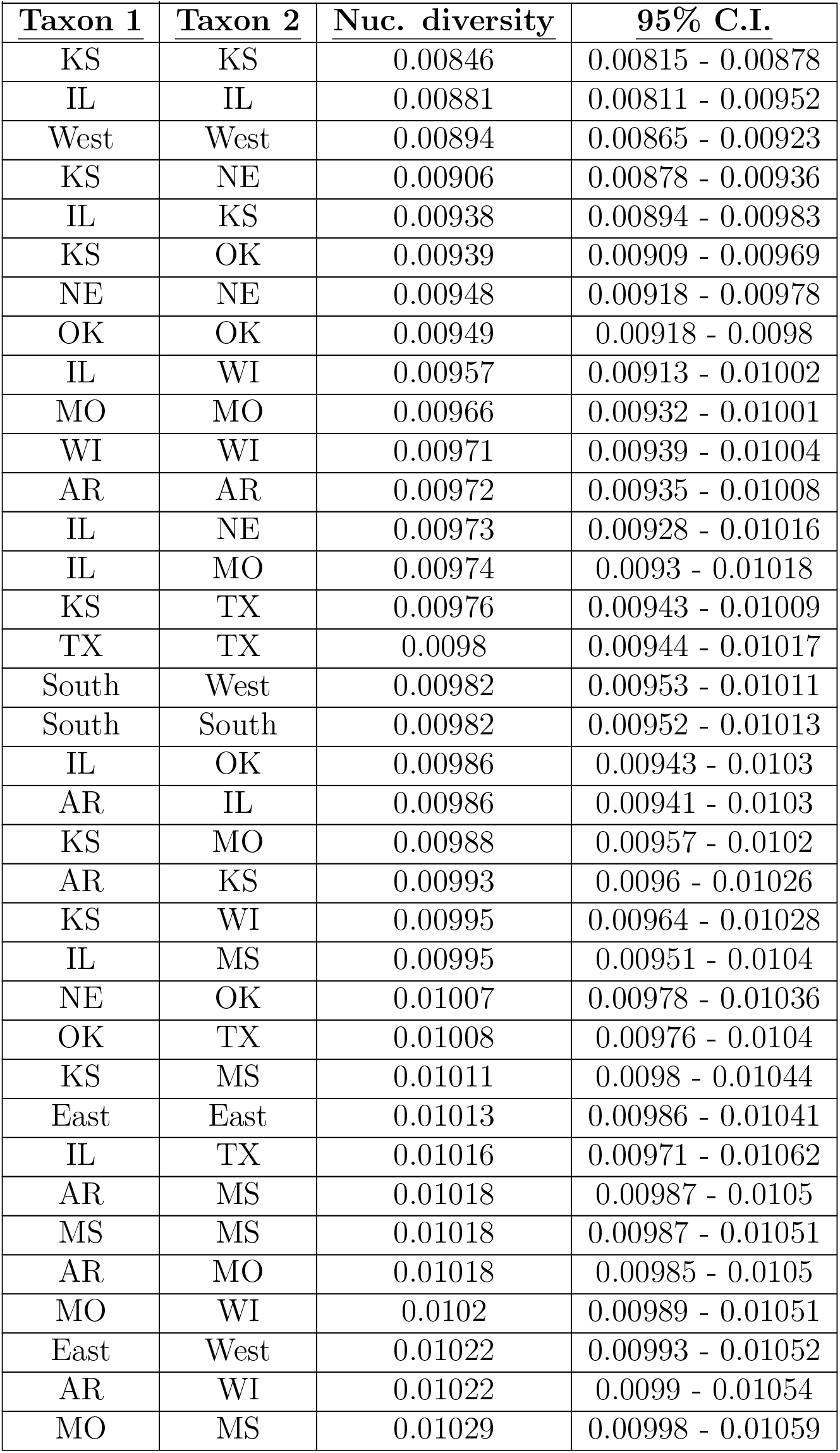
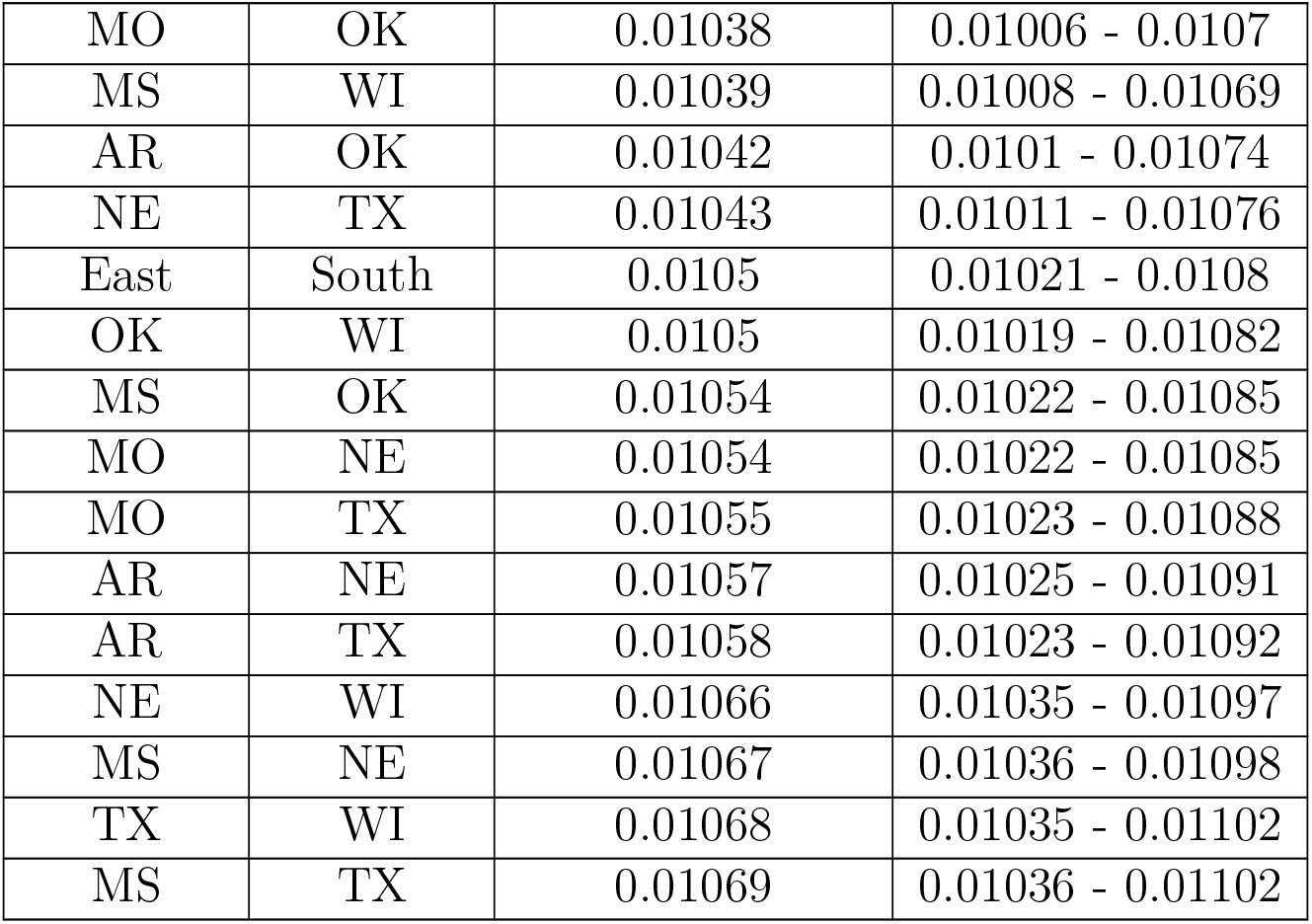
Summarized nucleotide variation found across {S. integrifolium} samples. Mean *π* (same Taxon 1 and Taxon 2) and *D_xy_* (different Taxon 1 and Taxon 2) and 95% confidence intervals, calculated from bootstrap replicates across contigs, are reported.

| <u>Taxon 1</u> | <u>Taxon 2</u> | <u>Nuc. diversity</u> | <u>95% C.I.</u> |
| --- | --- | --- | --- |
| KS | KS | 0.00846 | 0.00815 - 0.00878 |
| IL | IL | 0.00881 | 0.00811 - 0.00952 |
| West | West | 0.00894 | 0.00865 - 0.00923 |
| KS | NE | 0.00906 | 0.00878 - 0.00936 |
| IL | KS | 0.00938 | 0.00894 - 0.00983 |
| KS | OK | 0.00939 | 0.00909 - 0.00969 |
| NE | NE | 0.00948 | 0.00918 - 0.00978 |
| OK | OK | 0.00949 | 0.00918 - 0.0098 |
| IL | WI | 0.00957 | 0.00913 - 0.01002 |
| MO | MO | 0.00966 | 0.00932 - 0.01001 |
| WI | WI | 0.00971 | 0.00939 - 0.01004 |
| AR | AR | 0.00972 | 0.00935 - 0.01008 |
| IL | NE | 0.00973 | 0.00928 - 0.01016 |
| IL | MO | 0.00974 | 0.0093 - 0.01018 |
| KS | TX | 0.00976 | 0.00943 - 0.01009 |
| TX | TX | 0.0098 | 0.00944 - 0.01017 |
| South | West | 0.00982 | 0.00953 - 0.01011 |
| South | South | 0.00982 | 0.00952 - 0.01013 |
| IL | OK | 0.00986 | 0.00943 - 0.0103 |
| AR | IL | 0.00986 | 0.00941 - 0.0103 |
| KS | MO | 0.00988 | 0.00957 - 0.0102 |
| AR | KS | 0.00993 | 0.0096 - 0.01026 |
| KS | WI | 0.00995 | 0.00964 - 0.01028 |
| IL | MS | 0.00995 | 0.00951 - 0.0104 |
| NE | OK | 0.01007 | 0.00978 - 0.01036 |
| OK | TX | 0.01008 | 0.00976 - 0.0104 |
| KS | MS | 0.01011 | 0.0098 - 0.01044 |
| East | East | 0.01013 | 0.00986 - 0.01041 |
| IL | TX | 0.01016 | 0.00971 - 0.01062 |
| AR | MS | 0.01018 | 0.00987 - 0.0105 |
| MS | MS | 0.01018 | 0.00987 - 0.01051 |
| AR | MO | 0.01018 | 0.00985 - 0.0105 |
| MO | WI | 0.0102 | 0.00989 - 0.01051 |
| East | West | 0.01022 | 0.00993 - 0.01052 |
| AR | WI | 0.01022 | 0.0099 - 0.01054 |
| MO | MS | 0.01029 | 0.00998 - 0.01059 |
| MO | OK | 0.01038 | 0.01006 - 0.0107 |
| MS | WI | 0.01039 | 0.01008 - 0.01069 |
| AR | OK | 0.01042 | 0.0101 - 0.01074 |
| NE | TX | 0.01043 | 0.01011 - 0.01076 |
| East | South | 0.0105 | 0.01021 - 0.0108 |
| OK | WI | 0.0105 | 0.01019 - 0.01082 |
| MS | OK | 0.01054 | 0.01022 - 0.01085 |
| MO | NE | 0.01054 | 0.01022 - 0.01085 |
| MO | TX | 0.01055 | 0.01023 - 0.01088 |
| AR | NE | 0.01057 | 0.01025 - 0.01091 |
| AR | TX | 0.01058 | 0.01023 - 0.01092 |
| NE | WI | 0.01066 | 0.01035 - 0.01097 |
| MS | NE | 0.01067 | 0.01036 - 0.01098 |
| TX | WI | 0.01068 | 0.01035 - 0.01102 |
| MS | TX | 0.01069 | 0.01036 - 0.01102 |

**Appendix 3:** Description of climatic variables used in BayEnv2 analyses of genotype-environment associations.

| <u>Variable</u> | <u>Description</u> |
| --- | --- |
| EMT | extreme minimum temperature over 30 years |
| DD<18 | degree-days below 18C; heating degree-days |
| MAT | mean annual temperture |
| NFFD | number frost free days |
| MCMT | mean coldest month temperature |
| bFFP | the day of the year on which frost free period begins |
| FFP | frost free period |
| DD>5 | degree-days above 5C; growing degree-days |
| eFFP | the day of the year on which frost free period ends |
| Lat | latitude |
| DD<0 | degree days below 0C; chilling degree-days |
| PAS | precipitation as snow between August in previous year and July in current year |
| DD>18 | degree days above 18C; cooling degree days |
| Eref | Hargreaves reference evaporation |
| MWMT | mean warmest month temperature |
| TD | temperature difference between MWMT and MCMT (continentality) |
| MAR | mean annual solar radiation |
| AHM | annual heat-moisture index |
| SHM | summer heat-moisture index |
| EXT | extreme maximum temperature over 30 years |
| MAP | mean annual precipitation |
| long | longitude |
| CMD | Hargreaves climatic moisture deficit |
| MSP | May to September precipitation |
| RH | Relative Humidity |

**Appendix S4:**
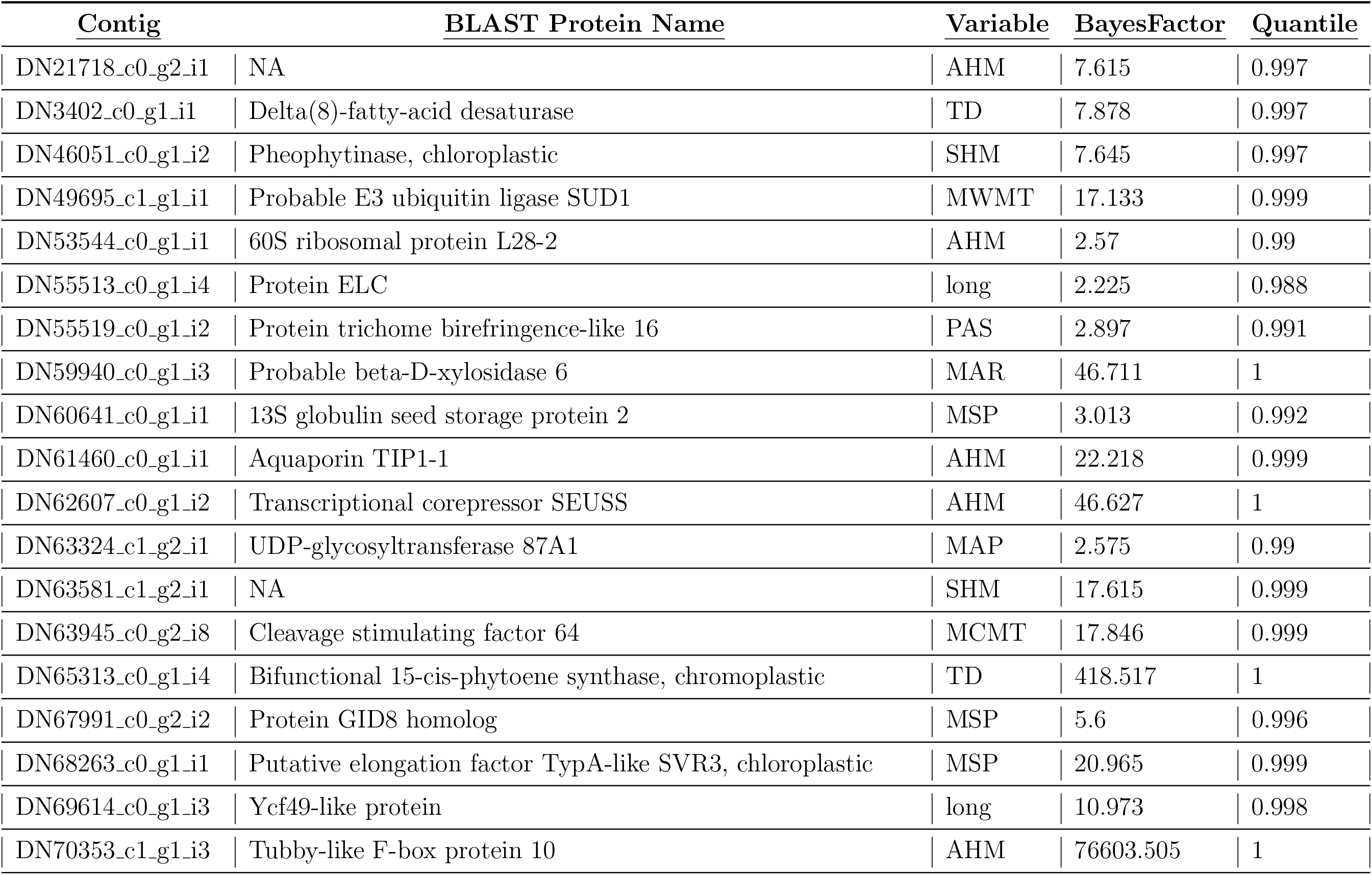
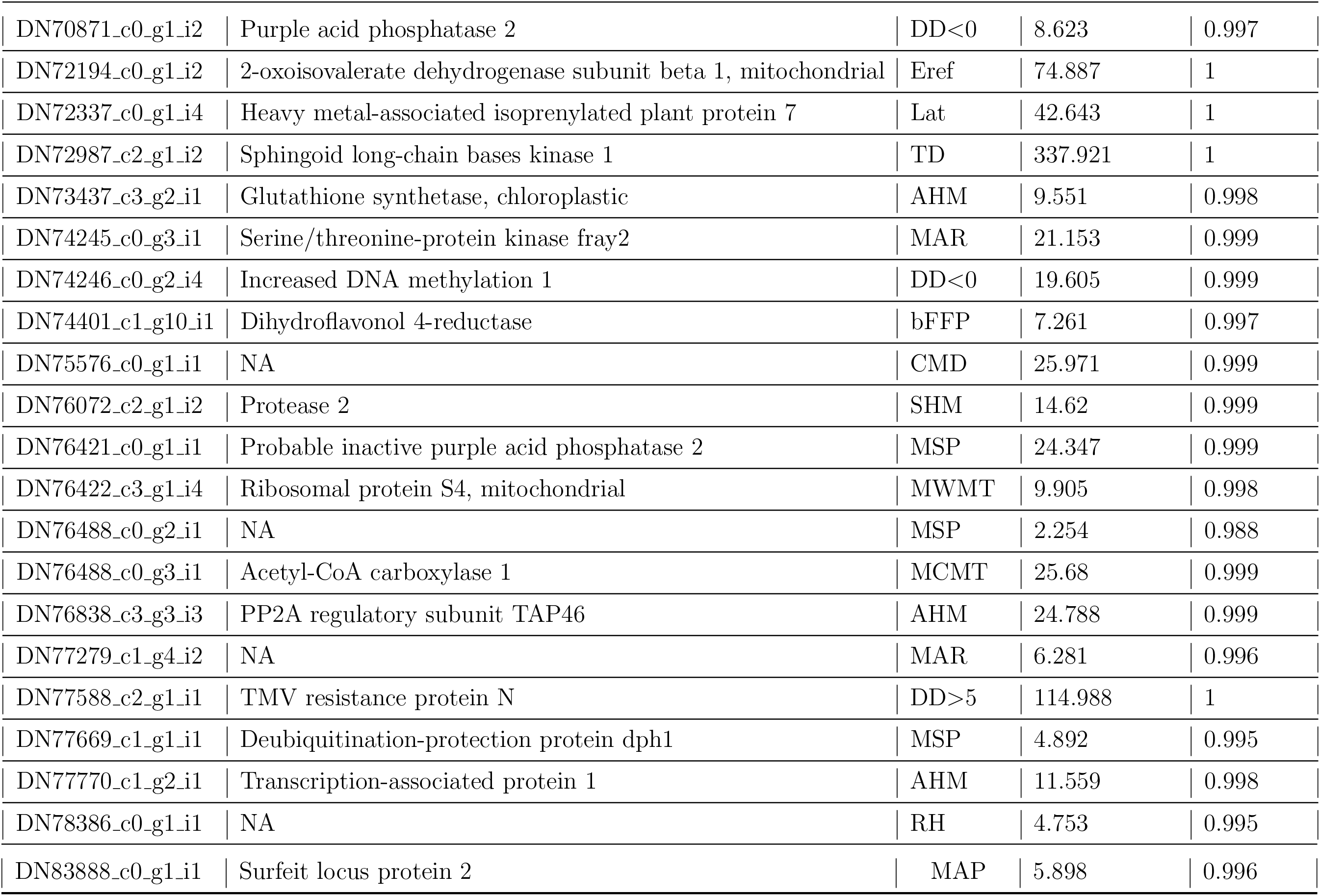
Associations between BayeScan inferred divergence outlier proteins and environmental variables sorted by highest quantile Bayes Factor. Environmental associations were estimated using BayEnv2.

**Appendix S5:** Associations between BayeScan inferred divergence outlier proteins and environmental variables sorted by highest quantile of absolute Pearson correlation coefficient. Environmental associations were estimated using BayEnv2.

| <u>Contig</u> | <u>BLAST Protein Name</u> | <u>Variable</u> | <u>Pearson</u> | <u>Quantile</u> |
| --- | --- | --- | --- | --- |
| DN21718_c0_g2.i1 | NA | PAS | 0.305 | 0.999 |
| DN3402_c0_g1.i1 | Delta(8)-fatty-acid desaturase | MAP | 0.237 | 0.994 |
| DN46051_c0_g1.i2 | Pheophytinase, chloroplastic | EXT | 0.23 | 0.993 |
| DN49695_c1_g1.i1 | Probable E3 ubiquitin ligase SUD1 | long | 0.332 | 0.999 |
| DN53544_c0_g1.i1 | 60S ribosomal protein L28-2 | DD>18 | 0.221 | 0.991 |
| DN55513_c0_g1.i4 | Protein ELC | AHM | 0.317 | 0.999 |
| DN55519_c0_g1.i2 | Protein trichome birefringence-like 16 | PAS | 0.229 | 0.993 |
| DN59940_c0_g1.i3 | Probable beta-D-xylosidase 6 | long | 0.399 | 1 |
| DN60641_c0_g1.i1 | 13S globulin seed storage protein 2 | long | 0.214 | 0.99 |
| DN61460_c0_g1.i1 | Aquaporin TIP1-1 | long | 0.459 | 1 |
| DN62607_c0_g1.i2 | Transcriptional corepressor SEUSS | MAR | 0.243 | 0.995 |
| DN63324_c1_g2.i1 | UDP-glycosyltransferase 87A1 | EXT | 0.155 | 0.962 |
| DN63581_c1_g2.i1 | NA | DD>18 | 0.274 | 0.998 |
| DN63945_c0_g2.i8 | Cleavage stimulating factor 64 | RH | 0.248 | 0.995 |
| DN65313_c0_g1.i4 | Bifunctional 15-cis-phytoene synthase, chromoplastic | NFFD | 0.327 | 0.999 |
| DN67991_c0_g2.i2 | Protein GID8 homolog | long | 0.278 | 0.998 |
| DN68263_c0_g1.i1 | Putative elongation factor TypA-like SVR3, chloroplastic | MAP | 0.211 | 0.989 |
| DN69614_c0_g1.i3 | Ycf49-like protein | AHM | 0.297 | 0.999 |
| DN70353_c1_g1.i3 | Tubby-like F-box protein 10 | long | 0.428 | 1 |

| Contig | BLAST Protein Name | Variable | Pearson | Quantile |
| --- | --- | --- | --- | --- |
| DN70871_c0_g1.i2 | Purple acid phosphatase 2 | MWMT | 0.328 | 0.999 |
| DN72194_c0_g1.i2 | 2-oxoisovalerate dehydrogenase subunit beta 1, mitochondrial | MSP | 0.227 | 0.992 |
| DN72337_c0_g1.i4 | Heavy metal-associated isoprenylated plant protein 7 | NFFD | 0.197 | 0.985 |
| DN72987_c2_g1.i2 | Sphingoid long-chain bases kinase 1 | FFP | 0.311 | 0.999 |
| DN73437_c3_g2.i1 | Glutathione synthetase, chloroplastic | long | 0.254 | 0.996 |
| DN74245_c0_g3.i1 | Serine/threonine-protein kinase fray2 | long | 0.292 | 0.998 |
| DN74246_c0_g2.i4 | Increased DNA methylation 1 | EXT | 0.16 | 0.967 |
| DN74401_c1_g10.i1 | Dihydroflavonol 4-reductase | AHM | 0.242 | 0.995 |
| DN75576_c0_g1.i1 | NA | TD | 0.296 | 0.999 |
| DN76072_c2_g1.i2 | Protease 2 | long | 0.281 | 0.998 |
| DN76421_c0_g1.i1 | Probable inactive purple acid phosphatase 2 | long | 0.223 | 0.992 |
| DN76422_c3_g1.i4 | Ribosomal protein S4, mitochondrial | long | 0.225 | 0.992 |
| DN76488_c0_g2.i1 | NA | MAR | 0.222 | 0.992 |
| DN76488_c0_g3.i1 | Acetyl-CoA carboxylase 1 | long | 0.368 | 1 |
| DN76838_c3_g3.i3 | PP2A regulatory subunit TAP46 | PAS | 0.273 | 0.997 |
| DN77279_c1_g4.i2 | NA | PAS | 0.254 | 0.996 |
| DN77588_c2_g1.i1 | TMV resistance protein N | DD<18 | 0.186 | 0.981 |
| DN77669_c1_g1.i1 | Deubiquitination-protection protein dph1 | long | 0.217 | 0.991 |
| DN77770_c1_g2.i1 | Transcription-associated protein 1 | DD<0 | 0.218 | 0.991 |
| DN78386_c0_g1.i1 | NA | EXT | 0.283 | 0.998 |

| <u>Contig</u> | <u>BLAST Protein Name</u> | <u>Variable</u> | <u>Pearson</u> | <u>Quantile</u> |
| --- | --- | --- | --- | --- |
| DN83888_c0_g1_i1 | Surfeit locus protein 2 | MAR | 0.269 | 0.997 |

**Appendix S6:**
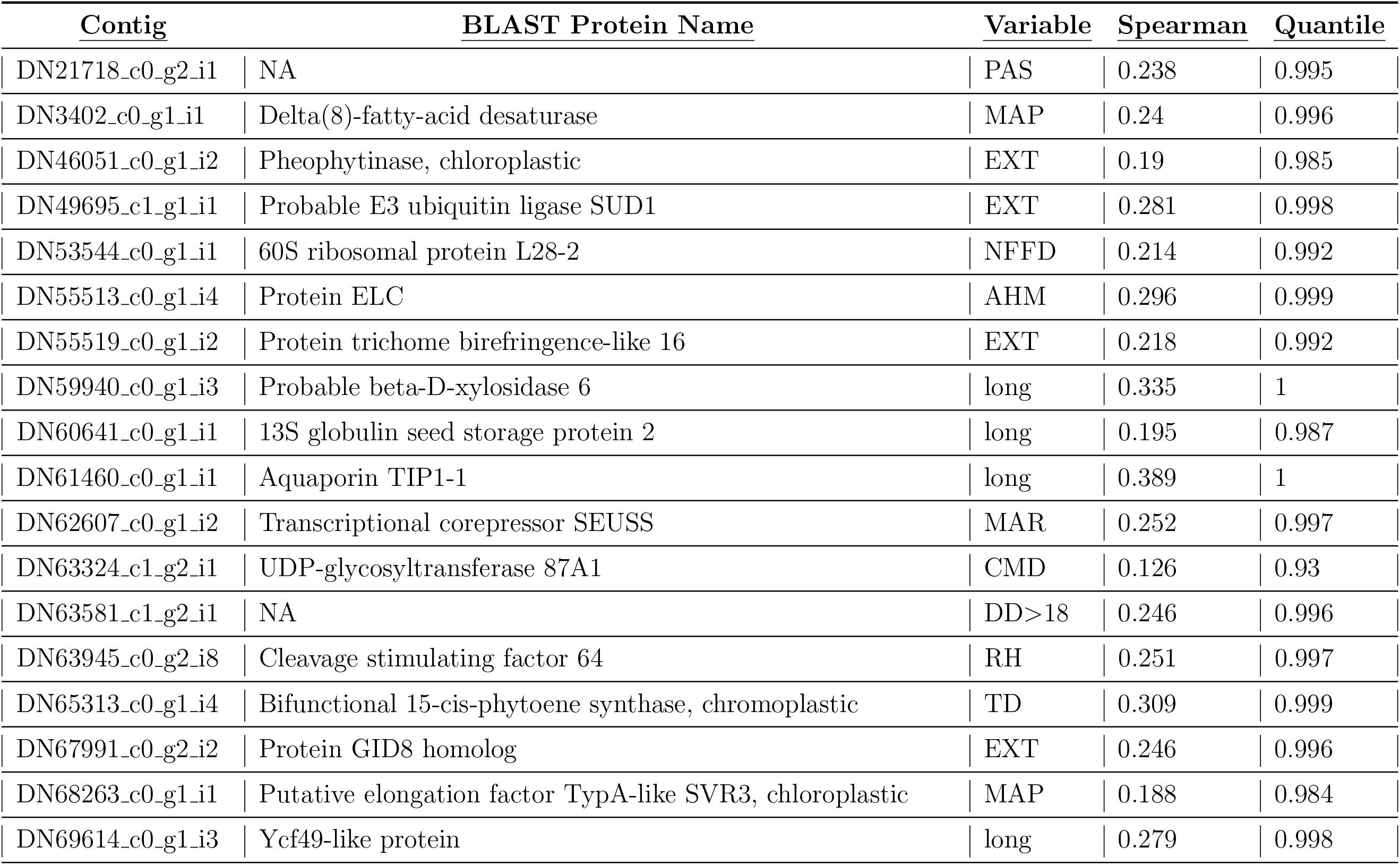
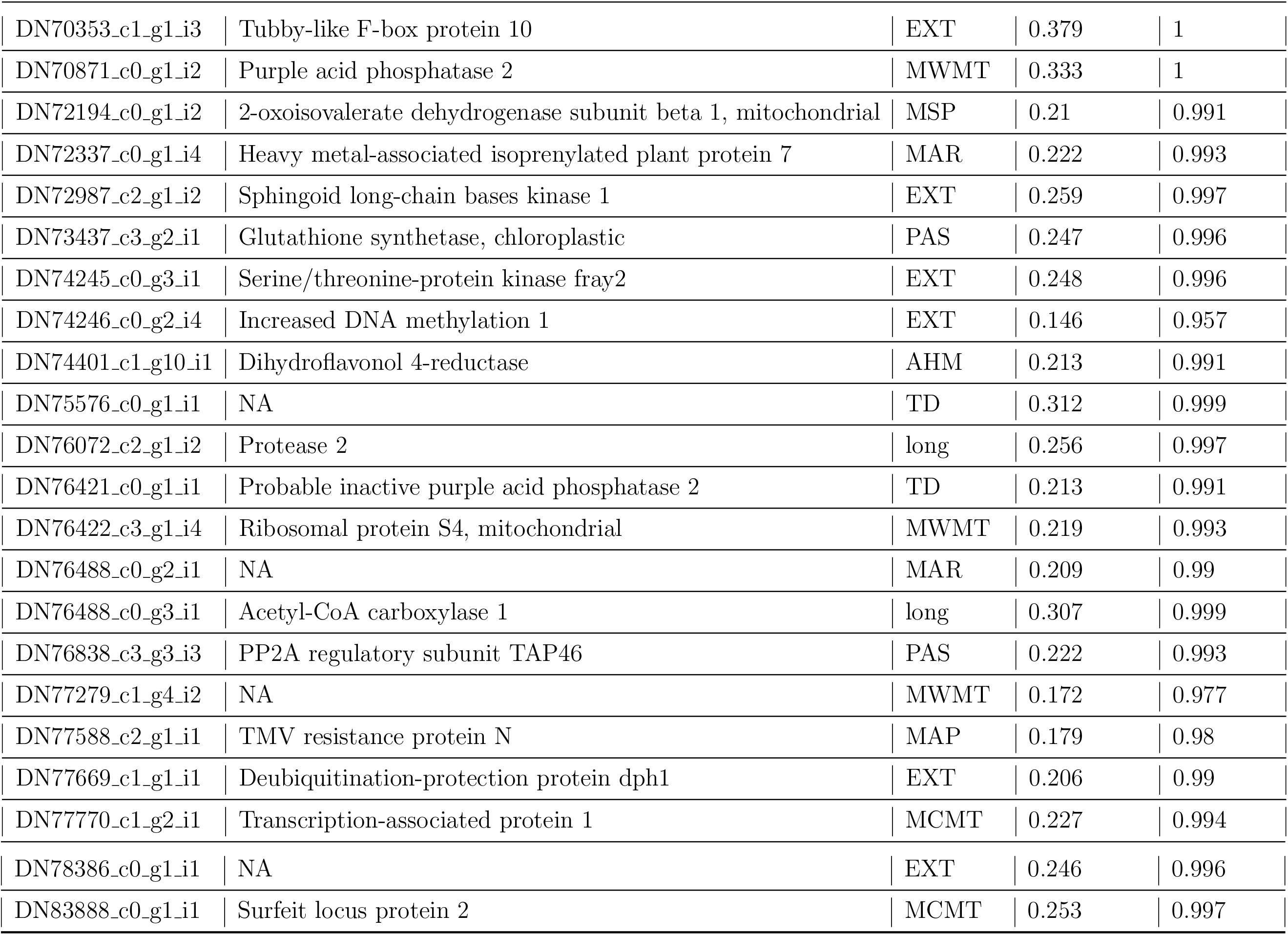
Associations between BayeScan inferred divergence outlier proteins and environmental variables sorted by highest quantile of absolute Spearman’s rank correlation coefficient. Environmental associations were estimated using BayEnv2.

**Appendix S7:**
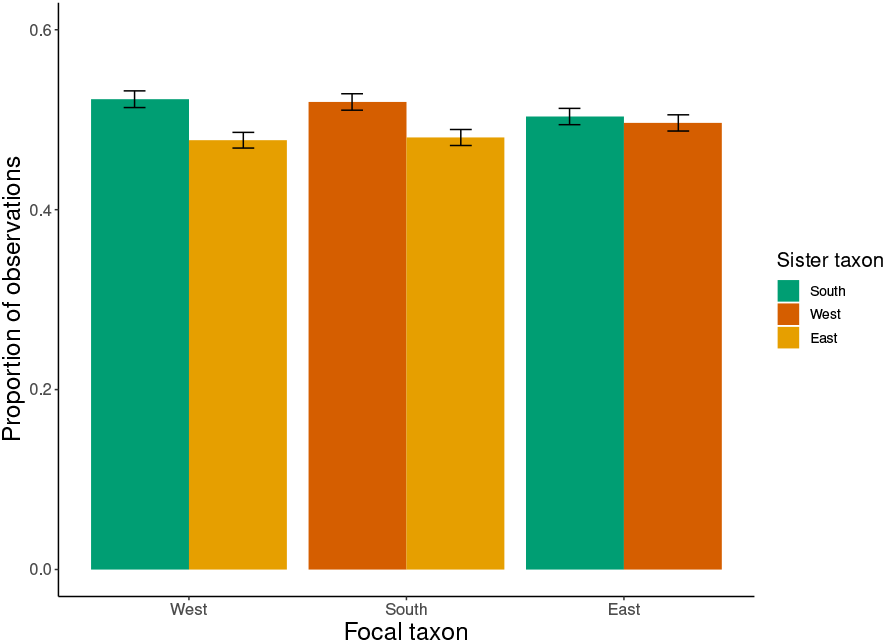
Summary of `TWISST` support for the `SVD Quartets` inferred phylogeny topology. The region of the sister tip for each focal taxon was recorded for each phylogeny subsampled in `TWISST` analysis across all marker contigs. Confidence around proportion estimates was estimated with 25,000 bootstrap replicates across contig phylogeny topologies. The significant excess of southern and western tips being found as sister supports their hypothesized most recent split (chi-square test, *p* < 0.05).

Appendix S8: Methods for MaxEnt species distribution modeling. We obtained *S. integrifolium* geo-referenced occurrence points from the Global Biodiversity Information Facility (GBIF) database. We then fit the maximum entropy species distribution model with five-fold cross validation using the `R` package `dismo`. Bioclimatic variables from the WorldClim database at 2.5 minutes of a degree resolution were used to train the model. Predicted habitat suitability was then mapped as an estimate of the species range.

